# p53Y220C-BET-bifunctionals (tPRIMEs) drive p53Y220C-mutant cancer cells into apoptosis

**DOI:** 10.1101/2025.08.13.669398

**Authors:** Charlotte Kelley, Jae Young Ahn, Scott Rusin, Thibaut Imberdis, Kong Xiong, Taylor Bramhall, Florence F. Wagner, Alexandra Joseph, Joshua Murtie, Moses Moustakim, Alexandra Lantermann

## Abstract

The transcription factor and tumor suppressor p53 is one of the most frequently mutated genes in cancer and has been difficult to target therapeutically due to its intrinsically disordered regions. The hotspot mutation p53Y220C, a mutation thermodynamically destabilizing p53, creates a unique extended crevice on the surface of the protein for which chemical matter has been identified over the last years. Advanced p53Y220C stabilizers reconstitute p53Y220C to its wildtype conformation, thereby restoring p53’s role in target gene expression and inhibiting the growth of p53Y220C mutant cancer cell lines. We hypothesized that direct recruitment of the transcriptional elongation machinery to p53Y220C and its target genes may potentiate effects beyond p53 protein stabilization alone. We leveraged induced proximity to discover bifunctional molecules, p53Y220C-**t**argeted **PR**oximity **I**nduced **M**odulators of **E**xpression (tPRIMEs), that specifically recognize the BET bromodomain proteins and induce stable ternary complexes with p53Y220C. p53Y220C-tPRIMEs potently inhibit proliferation and induce apoptosis of p53Y220C mutant cancer cell lines to a greater extent than the parental ligands alone or in combination. Gene expression analyses revealed that p53Y220C-tPRIMEs induce an increase in p53 target gene expression compared to parental binders. The superior antiproliferative activity, enhanced apoptosis, and increased p53 target gene expression are dependent on ternary complex formation. These data strongly suggest that a p53Y220C-tPRIME-mediated induced proximity approach between transcriptional regulators and p53Y220C - in contrast to p53 stabilization alone - can modulate the cell fate control from cell cycle inhibition to an apoptotic response, providing a compelling therapeutic modality for p53 mutant cancers.

## Materials and Methods

### Cell Culture

Gastric cancer cell line (NUGC3), hepatocellular cancer cell line (HUH7), oral cavity cancer cell line (KON), pancreatic cancer cell lines (BXPC3, T3M4), colorectal cancer cell line (HCT116) and lung cancer cell line (H1299) were obtained from American Type Culture Collection (ATCC) or Japanese Collection of Research Bioresources Cell Bank (JCRB). Cells were cultured in appropriate media (DMEM or RPMI-1640) supplemented with 10% FBS and incubated at 37 °C with 5% CO_2_ in a humidified atmosphere.

### CellTiter-Glo (CTG) cell proliferation assay

Cell viability was assessed by quantifying ATP using the CellTiter-Glo assay (Promega, #G9243). To determine the effect of compounds on cell viability, 2,000 to 5,000 cells/well were plated in 96-well plates using appropriate media and treated the following day with compounds at a 10-point dilution series for 96 hours. CTG reagent was then added, and luminescence signals were measured using a PHERAstar FSX Plate Reader (BMG Labtech). The percent viability was calculated using the following formula:

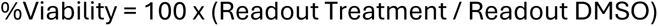

A four-parameter curve fit was employed using GraphPad PRISM to calculate IC_50_.

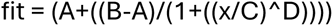

A: Bottom; B: Top; C: IC_50_; D: Hill Slope

### Cleaved Caspase3/7 apoptosis assay

Cleaved caspase 3/7 activity was monitored in cells by first seeding 1,250 cells per well of a solid white tissue culture treated 384-well plate (Corning, #3570) in 50 μl RPMI-1640+10% FBS or DMEM+10% FBS as appropriate for each cell line, then incubated overnight at 37 °C and 5% CO_2_. The following day, compounds were serially diluted in DMSO and then added to the cells at 1000x using an Echo 650 from an Echo LDV 384-well plate (#001-14615). Cells were incubated for 24 hours at 37 °C. To measure cleaved caspase 3/7 activity, 25 μl of Caspase-Glo 3/7 reagent (Promega, #G8090) were added to each well. Plates were shaken for 5 min at 400 rpm and incubated for another 10 minutes at room temperature. Luminescence signal was read out on a Pherastar plate reader (BMG Labtech). In parallel, an identical set of samples was measured for relative cell number by adding 25 μl of CTG reagent (Promega, #G9243) to each well, plates were then rocked at 400 rpm for 2 minutes, and luminescence signal was read out using a Pherastar plate reader (BMG Labtech). To analyze the data, cleaved caspase luminescence signal was normalized to CTG luminescence signal, then all values were normalized to DMSO control.

### Ternary complex ELISA assay

50,000 NUGC3 cells were plated in 96-well plates in RPMI-1640+10%FBS media and incubated overnight at 37 °C in 5% CO_2_. The next day, cells were treated with compounds for 4 hours at 37 °C. The media was then aspirated, and proteins were extracted by lysing the cells in lysis-buffer (Cell Signaling, #9803) supplemented with protease and phosphatase inhibitors (Thermo Fisher Scientific, #1861282). After diluting the lysate with ELISA Sample Diluent (Cell Signaling, #11083S) it was loaded onto an anti-p53 coated ELISA plate (Cell Signaling, #7370) and incubated overnight at 4 °C. The following day, the plate was washed in Phosphate Buffered Saline + 0.0025% Tween-20 (PBST) and then incubated with HRP-conjugated anti-BRD4 antibody (Cell Signaling, #77195CA) for 2 hours rocking at room temperature. After washing the plate in PBST, the plate was incubated with TMB substrate (Cell Signaling, #7004P) for 10 minutes at 37 °C. The reaction was then stopped with STOP solution (Cell Signaling, #7002P), and absorbance was measured at 450nm on the Glomax plate reader (Promega).

### Gene expression profiling by qPCR

30,000 NUGC3 cells were plated into 96-well plates in RPMI-1640+10%FBS media and incubated overnight at 37 °C in 5% CO_2_. The next day, cells were treated with compounds for 24 hours at 37 °C. Media was aspirated, and cells were lysed in buffer (50 mM Tris pH 8.0, 75 mM KCl, 6% Ficoll PM-400, 0.15% Triton X-100) containing protease and RNase inhibitors (Invitrogen, #10777019) suitable for one-step RT-PCR. Taqman reactions were set up using TaqMan™ Fast Virus 1-Step Master Mix (Thermo Fisher Scientific, #4444434), Taqman probes (Thermo Fisher Scientific), and lysate and run using a QuantStudio6. After an initial 5 min at 50 °C followed by 20 sec at 95 °C, samples underwent 38 cycles of 3 sec at 95 °C and 30 sec at 60 °C. CT values were quantified using the delta-delta CT method and results presented as fold change.

#### Probes

Probes:
| Gene Name | Probe ID (Thermo) |
| --- | --- |
| BBC3 | Hs00248075_m1 |
| PMAIP1 | Hs00560402_m1 |
| NQO1 | Hs05639726_s1 |
| ATF3 | Hs00231069_m1 |
| GAPDH | Hs99999905_m1 |
| TP53I3 | Hs00936519_m1 |
| CDKN1A | Hs01121172_m1 |

#### Reaction setup

Reaction setup:
| Volume (µl) | Reagent |
| --- | --- |
| 1.5 | 4x Taqman Master Mix |
| 0.3 | 20x probe for gene of interest |
| 0.1 | 60x housekeeping GAPDH |
| 1 | Cell Lysate |
| 3.1 | H<sub>2</sub>O |
| 6 | Total volume |

### SDS gel and Western Blot

Cell lysates were generated in RIPA lysis buffer (Thermo, #89901) or Pierce IP Lysis Buffer (Thermo Fisher Scientific, #87787) supplemented with protease and phosphatase inhibitors (Thermo Fisher Scientific, #1861282) and nuclease (Thermo Fisher Scientific, #88701). The lysate was precleared by centrifugation at 15,000 rpm at 4 °C for 10 min, and the protein concentration was quantified. 20μg of protein was separated by SDS-PAGE gel electrophoresis (Thermo Fisher Scientific, #WG1401BOX) and transferred to a nitrocellulose membrane (Biorad, #1704159). After blocking the membranes in Intercept Blocking Buffer (Licor, #927-6000), membranes were incubated overnight at 4 °C in different primary antibodies diluted 1/1000 in Intercept Antibody Diluent (Licor, 927-65001). The following list of primary antibodies was used: anti-p53 (SantaCruz, #sc-126), anti-p21 (SantaCruz, #sc-53870), anti-NQO1 (CST, #3187), anti-TP53I3 (Origene, #TA503656), anti-PUMA (CST, #12450), anti-cleaved Parp (CST, #5625), anti-Parp (CST, #9542), anti-MDM2 (CST, #86934), anti-BRD2 (CST, #5848), anti-BRD3 (CST, #94032), anti-BRD4 (CST, #13440), anti-GAPDH (CST, #97166), anti-β-actin (CST, #3700).

The following day, membranes were washed three times in TBST and then incubated in Donkey anti-Rabbit/anti-Mouse IR680 or IR800 antibody (Licor, #926-68073; #926-68072) diluted 1/5000 for 1 hour at room temperature. Membranes were washed three times in TBST and then imaged on the Azure imager.

### Immunoprecipitation

2 million NUGC3 cells were seeded into 10-cm dishes in RPMI-1640+10% FBS and incubated overnight at 37 °C and 5% CO_2_. The following day, cells were treated with compounds for 4 hours at 37 °C. Media was aspirated, and plates were washed twice in cold PBS. Next, protein was extracted by lysing the cells in Pierce IP Lysis Buffer (Thermo Fisher Scientific, #87787) supplemented with protease and phosphatase inhibitors (Thermo Fisher Scientific, #1861282) and nuclease (Thermo Fisher Scientific, #88701). The lysate was precleared by centrifugation at 15,000 rpm at 4 °C for 10min, and the protein concentration was quantified. For immunoprecipitation, 600μg lysate was incubated with either anti-BRD4 antibody (Cell Signaling, #13440) or anti-p53 antibody (SantaCruz, sc-126) rotating overnight at 4 °C. The following day, either protein A (Cytiva, #28-9440-06) or protein G (Cytiva, #28-9513-79) coated magnetic beads were incubated with the IP-lysates for 1 hour at room temperature, respectively. The beads were then washed with TBS + 0.0025% Tween-20 prior to being eluted by LDS sample buffer (Thermo Fisher Scientific, #NP0008). Both input and IPs were separated by SDS-PAGE gel electrophoresis (Thermo Fisher Scientific, #WG1401BOX) and transferred to a nitrocellulose membrane (Biorad, #1704159). After blocking the membranes in Intercept Blocking Buffer (Licor, #927-6000), membranes were incubated overnight at 4°C in one of the following detection antibodies diluted 1/1000 in Intercept Antibody Diluent (Licor, #927-65001): anti-p53 antibody (Santa Cruz, #sc-126), anti-BRD2 (CST, #5848), anti-BRD3 (CST, #94032), anti-BRD4 (CST, #13440). The following day, membranes were washed three times in TBST and then incubated in Donkey anti-Rabbit IR680 or IR800 antibody (Licor, #926-68073; #926-68072) diluted 1/5000 for 1 hour at room temperature. Membranes were washed three times in TBST and then imaged on the Azure imager. Quantification of Western Blot coimmunoprecipitation intensities was performed using ImageJ.

### In vivo NUGC3 xenograft study

NUGC3 gastric cancer cells were expanded in RPMI-1640+10%FBS media. 2 million NUGC3 cells + Matrigel (1:1) were implanted subcutaneously into the flanks of Balb/c nude mice. Dosing of animals was started 6 days post inoculation at a tumor volume of 160 mm^3^ per group. Tumor volumes were measured by caliper three times per week. Percent tumor growth inhibition was calculated as follows: %TGI = (1 - (change in tumor volume in treatment group / change in tumor volume in control group)) × 100.

### Details on study design

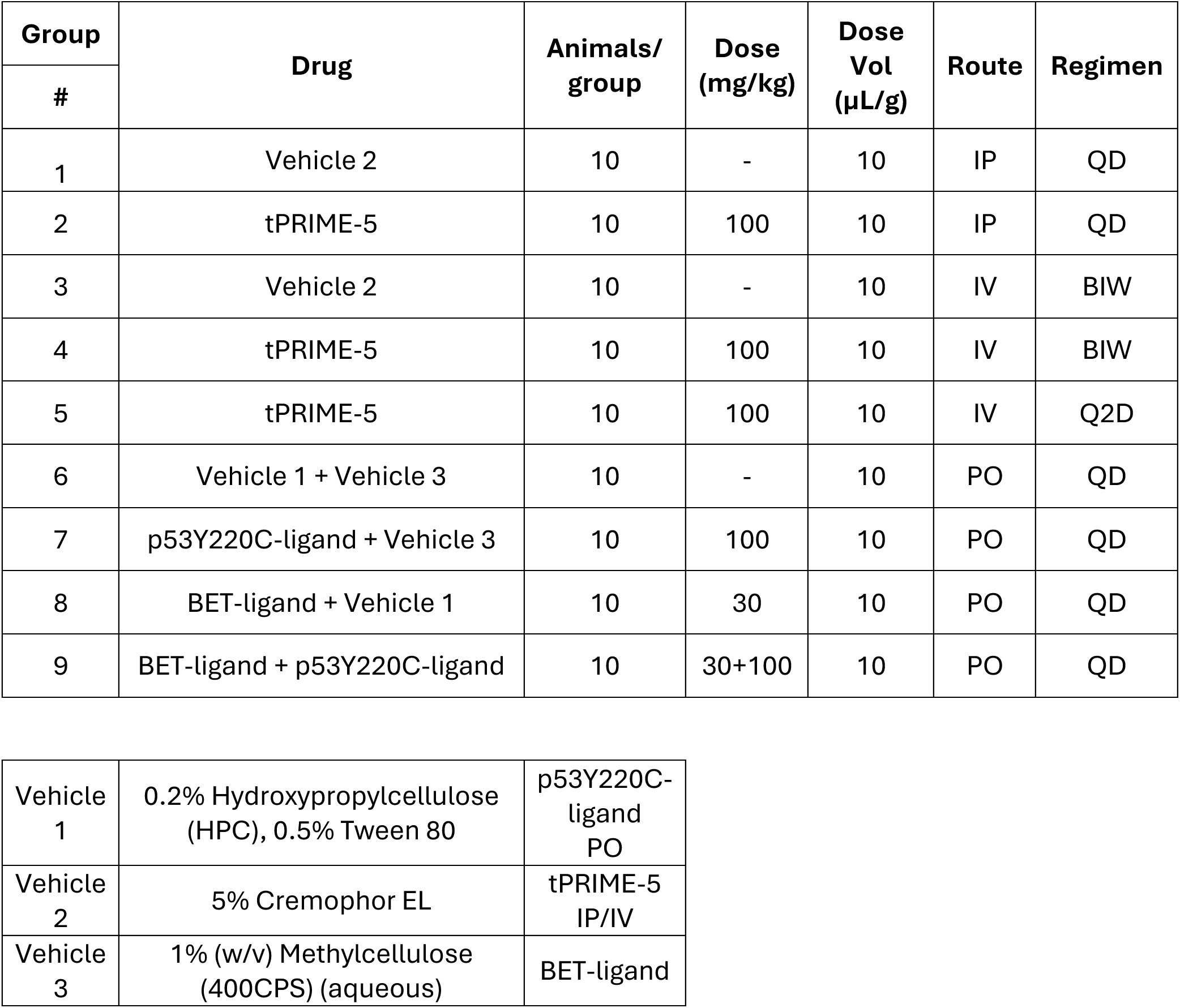

### Pharmacokinetic analysis of plasma and tumor samples

Serial concentrations of working solutions were prepared by diluting analyte stock solutions with blank plasma solution. 5 μl of working solutions (1, 2, 4, 10, 20, 100, 200, 1,000, 2,000ng/ml) were added to 10 μl of blank Balb/c nude mice plasma to achieve calibration standards of 0.5 ~1000 ng/ml (0.5, 1, 2, 5,10, 50, 100, 500, 1,000 ng/ml) in a total volume of 15 μl. Five quality control samples at 1 ng/ml, 2 ng/ml, 5 ng/ml, 50 ng/ml and 800 ng/ml for plasma were prepared independently of those used for the calibration curves. These QC samples were prepared on the day of analysis in the same way as calibration standards.

15 μl standards, 15 μl QC samples and 15 μl unknown samples (10 μl plasma with 5 μl blank solution) were added to 200 μl of acetonitrile containing IS mixture for precipitating protein respectively. Then the samples were vortexed for 30 s. After centrifugation at 4 °C, 4,000 rpm for 15 min, the supernatant was diluted 3 times with water. 20 μl of diluted supernatant was injected into the LC/MS/MS system for quantitative analysis.

For homogenization of the tumor samples, the tumor samples were mixed with water at a ratio of 1:6 (tumor weight (g) to water volume (ml)). The actual concentration (ng/ml) is the detected value (ng/g) multiplied by 7.

Serial concentrations of working solutions were prepared by diluting analyte stock solution with 50% acetonitrile in water solution. 15 μl of working solutions (1, 2, 4, 10, 20, 100, 200, 1,000 and 2,000 ng/ml) were added to 30 μl of blank Balb/c nude mice tumor (diluted with 6 folds of water) to achieve calibration standards of 0.5~1,000 ng/ml (0.5, 1, 2, 5,10, 50, 100, 500 and 1,000 ng/ml) in a total volume of 45 ml. Five quality control samples at 1 ng/ml, 2 ng/ml, 5 ng/ml, 50 ng/ml and 800 ng/ml for tumor were prepared independently of those used for the calibration curves. These QC samples were prepared on the day of analysis in the same way as calibration standards.

45 μl standards, 45 μl QC samples and 45 μl unknown samples (30 μl diluted tumor with 15 μl blank working solution) were added to 200 μl of acetonitrile containing IS mixture for precipitating protein respectively. Then the samples were vortexed for 30 s. After centrifugation at 4 °C, 4,000 rpm for 15 min, the supernatant was diluted 3 times with water. 5 μl of supernatant was injected into the LC/MS/MS system for quantitative analysis.

Concentrations of the compounds in the plasma and tumor tissue samples were analyzed using an LC-MS/MS method. WinNonlin (PhoenixTM, version 8.4) software was used for pharmacokinetic calculations. PK analyses of compounds were performed with AB API 6500+ LC/MS/MS instrument.

## Introduction

The tumor suppressor p53 is mutated in about 50% of human cancers, contributing to uncontrolled cell proliferation^1^. p53Y220C is a structural p53 mutation present in approximately 0.8% of human cancers^2^. This mutation gives rise to a crevice on the surface of p53Y220C, which thermally destabilizes the protein, leading to its aggregation and inactivation^3^. Ligands specifically binding to this surface crevice have been identified and optimized over the last years, and these p53Y220C-stabilizers have been shown to induce reactivation of p53 target genes and cell cycle arrest specifically in p53Y220C-mutant cells, providing an intriguing treatment option in precision oncology^2,4–7^. PMV Pharma’s molecule Rezatapopt is the most advanced p53 stabilizer and is currently in Phase 1/2 clinical trials for both liquid and solid tumors^6–8^.

The bromodomain and extraterminal (BET)-family proteins, consisting of BRD2, BRD3, BRD4 and BRD-T, act as epigenetic readers that regulate the expression of many genes associated with cell proliferation and immunity^9,10^. BET-proteins recognize acetylated lysines on histones and non-histone proteins through their bromodomain and interact with the positive transcription elongation factor b (P-TEFb) complex, thereby promoting transcription elongation through RNA polymerase II activity^11^. Given the relevance of BET-proteins for cancer growth, multiple inhibitors have been developed that potently bind the first (BD1) and/or second (BD2) bromodomain of BET-proteins^12^. While inhibition of BET proteins remains challenging in the clinic due to toxicity^13–15^, the large suite of BET ligands paired with their tight regulation of transcriptional machinery allows this family of proteins to be attractive for chemical modification of transcriptional signaling in cancer.

Chemically induced proximity through heterobifunctional small molecules has been an emerging field of ever-growing therapeutic modalities^16,17^: PROTACs for protein degradation^18,19^, PHICSs for phosphorylation^20,21^, DUBTACs for deubiquitination^22^, AceTACs for acetylation^23,24^, TRAMs for relocalization^25,26^, RIPTACs for proximity-induced inhibition of essential effector proteins^27,28^, and TCIPs/TRAPs for transcriptional activation^29,30^, just to name a few. TCIPs (Transcriptional/epigenetic Chemical Inducer of Proximity) targeting the transcriptional co-repressor BCL6 and chemically inducing proximity to BRD4 using JQ1 have been shown to de-repress BCL6, thereby promoting expression of BCL6-regulated apoptotic genes, potent cell growth inhibition and apoptosis in hematological cancer cell lines^29^. This concept has been expanded to BCL6-CDK9-TCIPs, using CDK9 as the effector recruiting P-TEFb to BCL6, similarly inducing apoptotic gene expression and cell growth inhibition^30^. Several heterobifunctional modalities have been applied to p53Y220C, including AceTACs, TRAPs and RIPTACs: p53Y220C-AceTACs demonstrated ternary complex formation and successful acetylation of p53, but further optimization is needed to achieve potent antiproliferative activity^23,24^. p53Y2220C-TRAPs promoted ternary complex formation between p53Y220C and BRD4 and induce the p53 target genes p21 and MDM2, but they currently lack induction of p53-induced apoptotic genes such as PUMA and potent proliferation inhibition^31^. Halo-PLK1-RIPTAC molecules induced proximity between the Halo-tag fused to p53Y220C and the essential protein PLK1. Selective compound accumulation due to high protein abundance of mutant p53 enables effective cell killing through inhibition of the essential protein PLK1^28^, which is mechanistically different from the transcriptional reactivation of BCL6/p53 by TCIPs/TRAPs^29–31^.

Here, we developed p53Y220C-BET heterobifunctional molecules called tPRIMEs, which promote the interaction of stabilized p53Y220C with BET proteins. This induced proximity leads to increased recruitment of the associated transcriptional machinery to p53Y220C, thereby boosting transcription of p53 target genes and changing the fate of p53Y220C-mutant cells from cell cycle inhibition to an apoptotic response.

## Results

### p53Y220C-tPRIME molecules inhibit proliferation and induce apoptosis of p53Y220C-mutant cancer cells

To test the hypothesis that p53Y220C-BET-heterobifunctionals can outperform p53Y220C-stabilizers (Fig. 1A), we designed a matrix of heterobifunctional molecules linking p53Y220C-ligands with BET-ligands through a variety of linkers (Fig. 1B). These p53Y220C-tPRIME molecules were screened in a 96h CTG cell proliferation assay in NUGC3, a p53Y220C mutant gastric cancer cell line, and HCT116, a p53 wildtype colorectal cancer cell line. p53Y220C-tPRIMEs inhibited proliferation of p53Y220C mutant NUGC3 cells more potently and demonstrated a greater E_max_ than the p53Y220C stabilizer alone (Fig. 1C). In addition, tPRIME molecules showed only weak or no effect on the proliferation of p53 wildtype HCT116 cells, suggesting selectivity for cancer cell lines carrying p53Y220C mutation (Fig. 1C). tPRIME-5 was the most potent and selective molecule, and was not only superior to p53Y220C-ligand, but was also more potent and efficacious than the BET-ligand alone or the combination of p53Y220C-ligand and BET-ligand (Fig. 1D). Given the difference in achieved E_max_, we assessed if tPRIME molecules induced apoptosis. Treatment with p53Y220C-tPRIME-5, but not with individual ligands alone or in combination, induced apoptosis as measured by cleaved caspase 3/7 assay 24h post treatment selectively in NUGC3 cells over HCT-116 cells (Fig. 1E). Lastly, besides demonstrating profound activity in NUGC3 cells, tPRIME-5 showed superior antiproliferative and pro-apoptotic response across additional p53Y220C-mutant cell lines, demonstrating the broader translatability of this concept across p53Y220C-mutant cancers (Supplementary Fig. 1A and B, Supplementary Table 1). Taken together, tPRIME molecules can selectively and potently induce apoptosis in p53Y220C cell lines, which in turn drives a greater reduction in cell viability than stabilization of p53Y220C can achieve alone.

**Figure 1:**
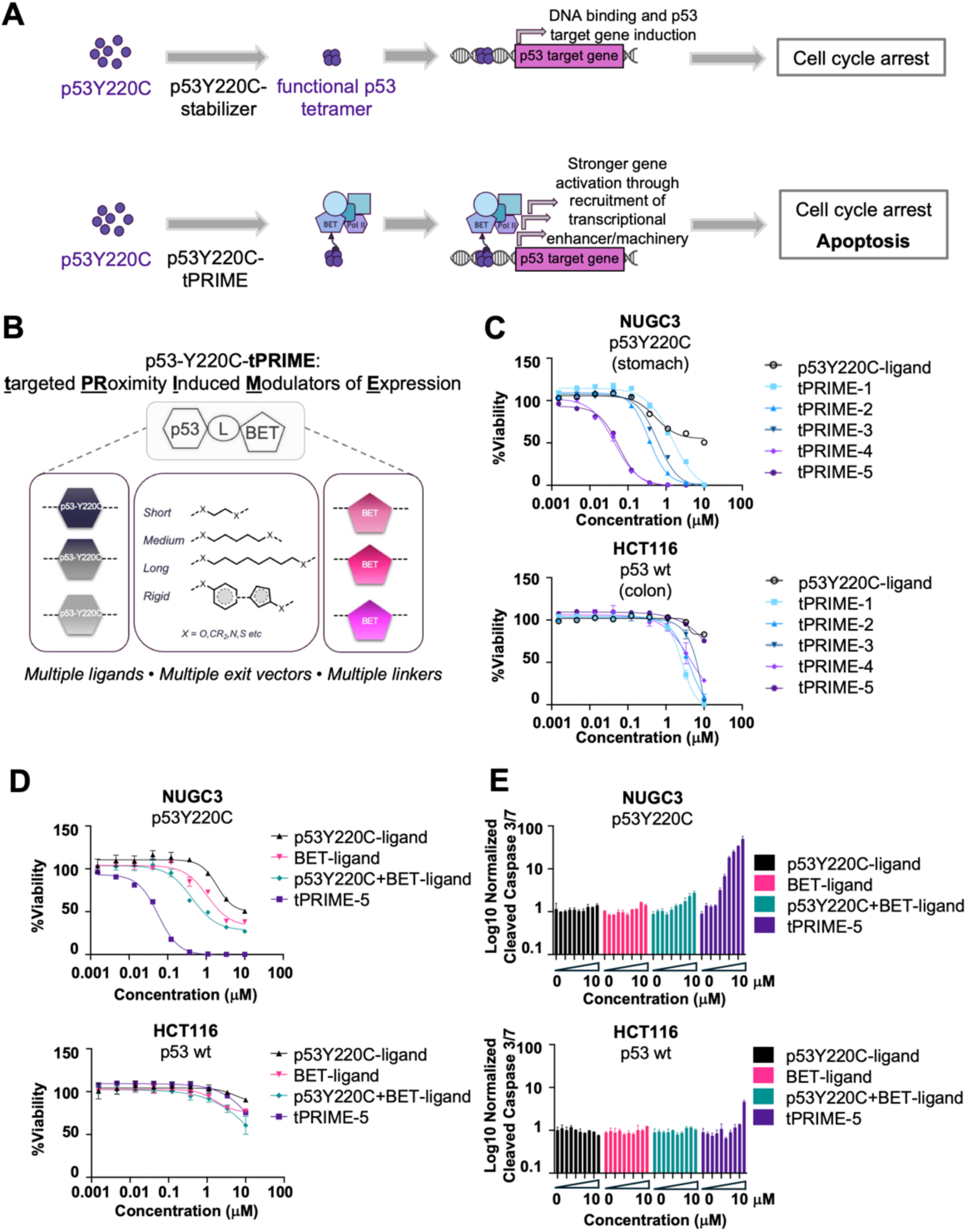
p53Y220C-tPRIMEs potently and selectively inhibit proliferation and induce apoptosis of NUGC3 p53Y220C-mutant cell line. **A.** Schematic showing the working hypothesis for p53Y220C-tPRIMEs versus p53Y220C-ligands. **B.** Illustration of matrix of tPRIMEs, linking different p53Y220C- and BET-ligands through a variety of linkers. **C.** Cell viability assessment via CTG after treatment of NUGC3 or HCT116 cells with tPRIME-1-5 or p53Y220C-ligand for 96h. **D.** Cell viability assessment via CTG after treatment of NUGC3 or HCT116 cells with tPRIME-5, p53Y220C-ligand, BET-ligand or equimolar combination of p53Y220C+BET-ligand for 96h. **E.** Apoptosis assessment via cleaved caspase 3/7 after treatment of NUGC3 or HCT116 cells with tPRIME-5, p53Y220C-ligand, BET-ligand or combination of p53Y220C+BET-ligand for 24h.

### p53Y220C-tPRIME molecules induce ternary complex formation between p53Y220C and BET-proteins

The enhanced activity of the heterobifunctional molecule compared to the p53Y220C-ligand and BET-ligand combination suggested an induced proximity gain of function mechanism of action. p53 and BRD4 have been previously shown to interact^32,33^, and we hypothesized that tPRIMEs were operating by inducing or potentiating the formation of ternary complexes between p53Y220C and BET proteins.

To assess interaction between p53Y220C and BRD4 biochemically, a p53Y220C-TR-FRET assay in the presence or absence of BRD4 was developed. In the absence of BRD4, tPRIME-5 showed lower binding affinity to p53Y220C than the p53Y220C-ligand alone (Fig. 2A, black curves). However, addition of BRD4-BD1-protein to the reaction profoundly increased the binding of tPRIME-5 to p53Y220C as reflected by the marked FRET signal loss due to probe displacement (Fig. 2A, top, pink curve). In contrast, the binding affinity of the p53Y220C-stabilizer to p53Y220C was not affected by the addition of BRD4 BD1-protein to the reaction (Fig. 2A, bottom, pink curve). These data demonstrated that tPRIME-5 induces a strong interaction between p53Y220C and BRD4 protein, which was attenuated by addition of excess of BET-ligand (Fig. 2A, blue curve). Likewise, endogenously in cells, antibody pulldown of BRD4 allowed the co-precipitation of p53 in cell lysates of NUGC3 cells treated with tPRIME-5, but not with p53Y220C- or BET-ligands alone or combined (Fig. 2B). Vice versa, pulldown of p53 following tPRIME-5 treatment allowed the co-precipitation of BRD4, as well as the two BET-domain containing proteins BRD2 and BRD3 (Supplementary Fig. 2A), and pulldown of BRD2 or BRD3 resulted in co-precipitation of p53 in a tPRIME-5 dependent manner (Supplementary Fig. 2F). Last, tPRIME-5-induced ternary complex formation was also detectable across a panel of additional p53Y220C-mutant cancer cell lines (Supplementary Fig. 2A-E). The formation of cellular tPRIME-induced ternary complex could also be confirmed more quantitatively through an ELISA assay based on a p53 capturing antibody and a BRD4 detection antibody (Fig. 2C). Together, these data demonstrate that p53Y220C-tPRIME molecules potently and reversibly induce a stable ternary complex between p53Y220C and BET-proteins.

**Figure 2:**
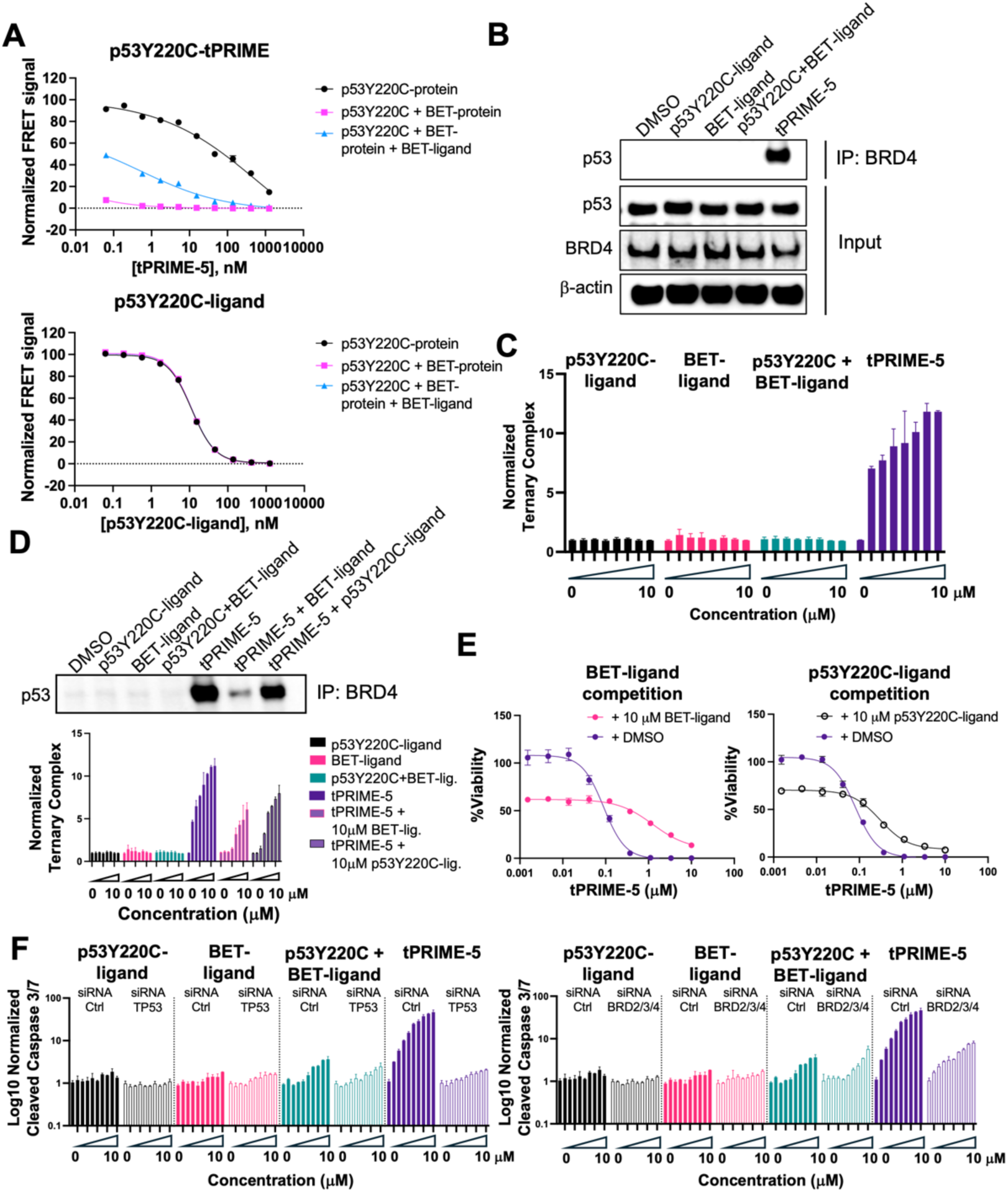
p53Y220C-tPRIMEs induce functionally relevant ternary complex. **A.** Assessment of binding of tPRIME-5 or p53Y220C-ligand to p53Y220C-protein in the presence or absence of BRD4-BD1-protein (+/- BET-ligand) via biochemical TR-FRET assay. **B.** Western blot demonstrating co-precipitation of p53 after pulldown of BRD4 in NUGC3 cells treated with 1μM tPRIME-5 or control ligands for 4h. **C.** ELISA measuring ternary complex formation after 4h treatment of NUGC3 cells with serially diluted tPRIME-5 or control ligands. **D.** ELISA measuring ternary complex formation after 4h treatment of NUGC3 cells with serially diluted tPRIME-5 in the presence or absence of 10μM BET-ligand or p53Y220C-ligand. **E.** 96h CTG cell proliferation assay of NUGC3 cells treated with tPRIME-5 in the presence or absence of 10μM BET-ligand or p53Y220C-ligand. **F.** Assessment of apoptosis via cleaved caspase3/7 in NUGC3 cells treated for 24h with tPRIME-5 or control ligands after knockdown of TP53 (p53) or BRD2/3/4.

### p53Y220C-tPRIME-induced ternary complex is required to inhibit cell proliferation and to induce apoptosis

To test if the formation of tPRIME-induced ternary complex between p53Y220C and BET-protein is required for tPRIME’s antiproliferative and pro-apoptotic function, chemical competition experiments were performed. Adding an excess of p53Y220C-ligand or BET-ligand to tPRIME-5 treated NUGC3 cells reduced ternary complex formation (Fig. 2D) and inhibited the activity of tPRIME-5 in a 96h CTG cell proliferation assay, as demonstrated by decreased potency and less pronounced E_max_ (Fig. 2E). In addition, ternary complex formation as measured by ELISA correlated with the potency of tPRIME molecules measured by CTG proliferation assay (Supplementary Fig. 3). Furthermore, genetic knockdown of p53 through siRNA prevented tPRIME-5-induced apoptosis in NUGC3 cells, and combined knockdown of BRD2, BRD3, and BRD4 also attenuated tPRIME-5-induced apoptosis (Fig. 2F). In contrast, individual knockdown of BRD2, BRD3, or BRD4 was not sufficient to prevent tPRIME-5-induced apoptosis, suggesting that all BET-proteins contribute to tPRIME’s cellular effects (Supplementary Fig. 4), consistent with reports of functional redundancy between the BET proteins.

### p53Y220C-tPRIME molecules induce transcription of p53 target genes associated with apoptosis induction

To elucidate the molecular mechanism of tPRIMEs, RNA-sequencing was performed of NUGC3 cells treated for 24h with two earlier stage tPRIME molecules, tPRIME-2 and tPRIME-3, or the p53Y220C- and BET control ligands. Similarly to tPRIME-5, the two earlier stage tool compounds tPRIME-2 and tPRIME-3 demonstrated more effective and more potent growth inhibition of p53Y220C-mutant cells than the p53Y220C-ligand alone (Fig. 1C). Hierarchical cluster analysis revealed a cluster of genes markedly upregulated by tPRIMEs compared to the control ligands at 24h (Fig. 3A). A pathway analysis of all genes upregulated by tPRIMEs using ENRICHR identified p53 signaling pathway to be more significantly modulated by tPRIMEs than by control ligands (Supplementary Fig. 5A and B). Similarly, a pathway analysis of the cluster of markedly upregulated genes identified through hierarchical clustering also revealed p53 target genes as the most prominently modulated pathway (Fig. 3A Table). The striking activation of p53 target genes led to a strong downregulation of cell cycle, DNA replication and E2F target genes, in alignment with p53’s role in activating the DREAM complex, a transcriptional repressor inhibiting cell cycle progression, DNA synthesis, G2/M transition and mitosis (Supplementary Fig. 5C)^34^. To evaluate the expression changes of upregulated genes in more detail, and to identify suitable biomarkers correlating with tPRIME’s phenotypic outcomes, gene expression of individual bona-fide p53 target genes^35^ was assessed using quantitative PCR or Western Blot. p53 target genes associated with cell cycle arrest, such as CDKN1A/p21 and MDM2, were induced more potently, but to a lower extent, by tPRIMEs compared to p53Y220C-stabilizer (Fig. 3B). In contrast, p53 target genes associated with the induction of apoptosis, such as BBC3/PUMA, ATF3 and TP53I3, were stimulated more potently and to a much greater extent by tPRIMEs than by p53Y220C-stabilizers (Fig. 3B). Encouragingly, tPRIME-5 induced expression of these genes to a greater extent than tPRIME-3, correlating with the superior anti-proliferative and pro-apoptotic potencies. Interestingly, the equimolar combination treatment with p53Y220C- and BET-ligand impaired p53Y220C-ligand mediated expression of all p53 target genes assessed (Fig. 3B).

**Figure 3:**
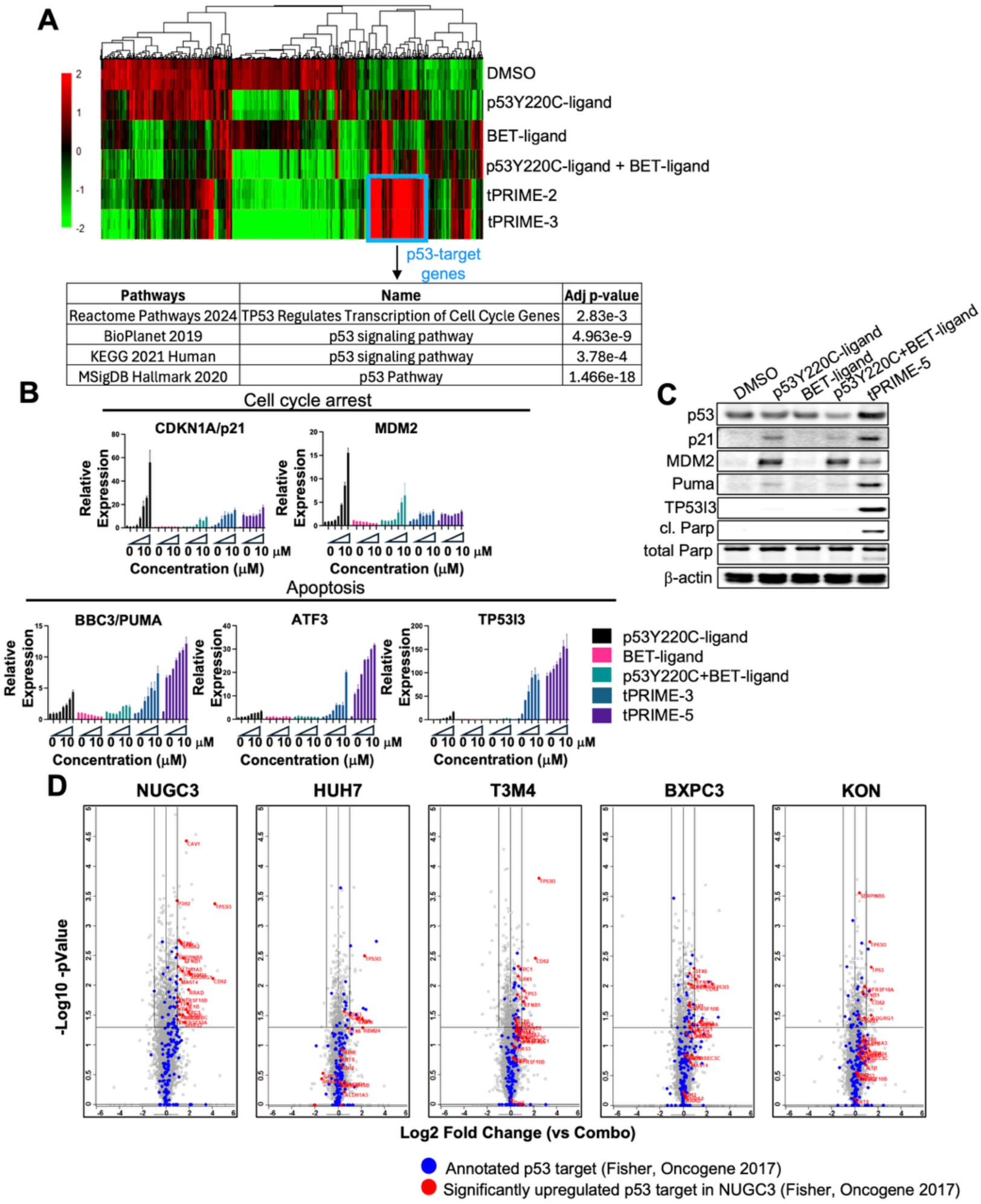
p53Y220C-tPRIMEs markedly induce p53 target genes. **A.** Hierarchical cluster analysis of genes identified as significantly upregulated in RNA-sequencing by 24h treatment with tPRIME-2, tPRIME-3 or control ligands compared to DMSO. Pathway analysis of cluster of genes significantly upregulated by tPRIME-2/3 compared to control ligands. **B.** qPCR analysis of the p53 target genes CDKN1A/p21, MDM2, BBC3, ATF3, TP53I3 in NUGC3 cells treated with serial dilution of tPRIME-3, tPRIME-5 or control ligands. **C.** Western Blot analysis of p53-dependent and apoptotic proteins in NUGC3 cells treated for 24h with 1μM of tPRIME-5 or control ligands. **D.** LC-MS/MS analysis of proteomic changes across multiple p53Y220C-mutant cell lines treated for 24h with 1μM of tPRIME-5 compared to p53Y220C+BET-ligand combination.

These changes on the RNA level translated to the protein level as measured for several p53-dependent target proteins via Western Blot (Fig. 3C), and proteome-wide via LC-MS/MS (Supplementary Fig. 6A and B; Fig 3D). Also, here, the more efficacious tPRIME-5 demonstrated more potent induction of p53 dependent target proteins than tPRIME-3 (Supplementary Fig. 6A and B). tPRIME-5 also upregulated p53 dependent target proteins to a greater extent than control treatments across additional p53Y220C-mutant cell lines, demonstrating the translation of the molecular mechanism across p53Y220C-mutant cancer cell lines (Fig. 3D).

We also assessed whether tPRIME-5 induces a more pronounced deregulation of genes modulated by BET-ligands. This would be analogous to the molecular mechanism described for RIPTAC molecules^27,28^, where selective cell proliferation inhibition and induction of apoptosis would result from BET-protein inhibition, and p53Y220C would serve to sequester the heterobifunctional molecules within p53Y220C expressing cell. Analysis of LC-MS/MS data across a panel of p53Y220C-mutant cancer cell lines did not identify BET-ligand target proteins to be further down- or upregulated by tPRIME-5 than by BET-ligands, excluding a RIPTAC effect as the dominant molecular mechanism underlying cell killing (Supplementary Fig. 7).

Together, these data demonstrate that p53Y220C-tPRIMEs, by inducing proximity between p53Y220C and the BET proteins, can markedly induce transcription of pro-apoptotic p53 target genes across a panel of p53Y220C-mutant cell lines.

### p53Y220C-tPRIME molecules prevent induction of MDM2-mediated negative feedback loop

MDM2 is an E3-ubiquitin ligase that is directly transcriptionally activated by p53. In turn, MDM2 ubiquitinates p53 leading to its degradation, thereby initiating a negative feedback loop^36,37^. p53Y220C-tPRIMEs induced MDM2 gene and protein expression to a lesser extent but more potently compared to p53Y220C-stabilizers (Fig. 3B and C). The lack of MDM2 induction correlated with p53 total levels remaining constant in the presence of tPRIME-5 treatment, whereas p53Y220C-stabilizers strongly induced MDM2 expression and downregulation of p53 protein levels (Fig. 3C).

### p53Y220C-tPRIMEs demonstrate delayed p53 tetramer formation and slower kinetics towards p53 target gene induction compared to p53Y220C-stabilizers

The RNA-sequencing and LC-MS/MS proteomics data demonstrated marked induction of p53 target genes in NUGC3 cells treated with tPRIMEs for 24h. To explore the kinetics of gene induction in more detail, expression changes of several p53 target genes were assessed over a time course of 2 to 24 hours. Interestingly, tPRIMEs induced expression across multiple p53 target genes more slowly than the p53Y220C-stabilizer (Fig. 4A). For example, expression of BBC3, ATF3 and TP53I3 continued to increase over the measured time course of 24h after treatment with tPRIME-5, whereas expression of these genes peaked at 2-8h post treatment with p53Y220C-ligand (Fig. 4A). Similarly, for CDKN1A and MDM2, 1μM p53Y220C-ligand induced maximal gene induction at 4h post treatment followed by a rapid drop, whereas tPRIME-5 demonstrated different potencies and kinetics towards these genes (Fig. 4A).

**Figure 4:**
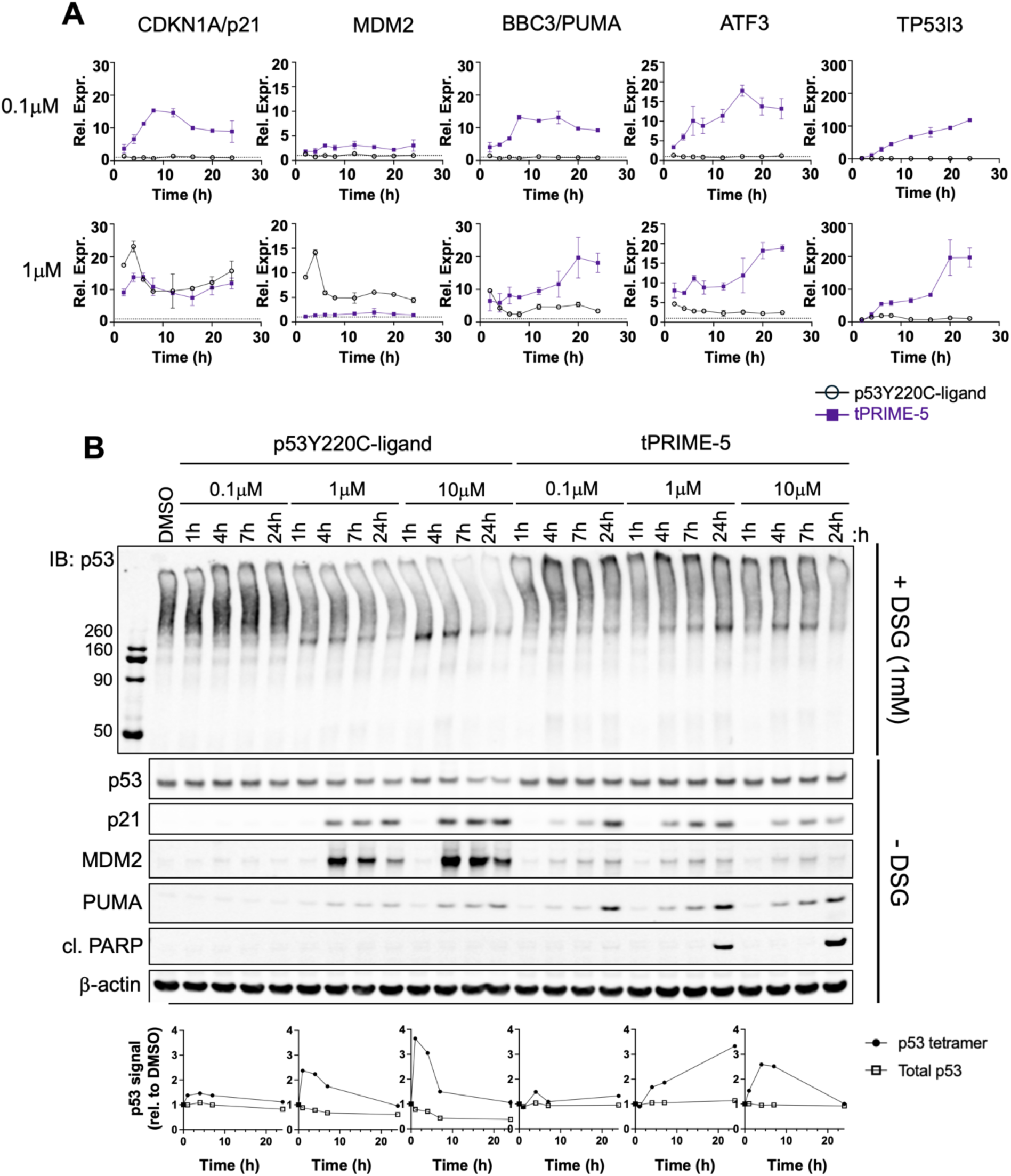
tPRIME-5 induces p53 target genes and tetramer formation at slower kinetics than p53Y220C-ligands. **A.** qPCR analysis of the p53 target genes CDKN1A, MDM2, BBC3, ATF3, TP53I3 in NUGC3 cells treated with 0.1 or 1μM of tPRIME-5 or p53Y220C-ligand over a time course of 2-24h. **B.** Western Blot analysis and quantification of p53 tetramer after DSG crosslinking in NUGC3 cells treated for 2-24h with 0.1μM, 1μM or 10μM p53Y220C-ligand or tPRIME-5.

Given that successful initiation of p53-dependent gene activation requires the formation of a p53 tetramer, we explored whether p53Y220C-tPRIME molecules induce tetramer formation to the same degree and with the same kinetics as the p53Y220C-ligand. The lysine-dependent crosslinker DSG was verified as an appropriate tool to capture the p53 tetramer (Supplementary Fig. 8A)^38^. In the presence of crosslinker, lysates from NUGC3 cells treated with DMSO or 0.1μM p53 Y220C-ligand appeared as smear at high molecular weight on the SDS gel, in agreement with the thermal instability of the p53Y220C mutant protein and tendency to aggregate. Treatment with 1μM or 10μM of p53Y220C-ligand induced tetramer formation, which was visible as a decrease in the high molecular weight smear and appearance of a distinct band around the expected p53 tetramer molecular weight of approximately 200kDa (Supplementary Fig. 8, Fig. 4B). This occurred quickly, with a peak at 1h post treatment and tetramer decreasing over the time course of 24h, correlating with the induction of MDM2 and decrease of p53 protein over time (Fig. 4B). In contrast, treatment with p53Y220C-tPRIME induced tetramer formation more potently than p53Y220C-stabilizer (~0.01μM), but with slower kinetics: barely any tetramer was visible at 1h while a gradual increase occurred over the time course of 24h (Fig. 4B, Supplementary Fig. 8B). As demonstrated above, treatment with tPRIME-5 barely induced MDM2 and as such p53 total levels remained constant (Fig. 4B).

In summary, these data demonstrate that p53Y220C-tPRIMEs induce tetramer formation more potently but at a slower kinetics compared to p53Y220C-stabilizers, correlating with the kinetics observed for induction of pro-apoptotic p53 target genes.

### p53Y220C-tPRIME molecules inhibit NUGC3 xenograft growth and lead to tumor regression

Analysis of the pharmacokinetic properties demonstrated poor oral bioavailability of tPRIME-5, but acceptably low clearance post intravenous (IV) and reasonable exposures after intraperitoneal (IP) dosing, covering the unadjusted cellular IC_50_ measured in NUGC3 (greater than ~2,500-fold coverage IV, greater than ~400-fold coverage IP based on C_0_ and C_max_, respectively) (Fig. 5A, Table).

**Figure 5:**
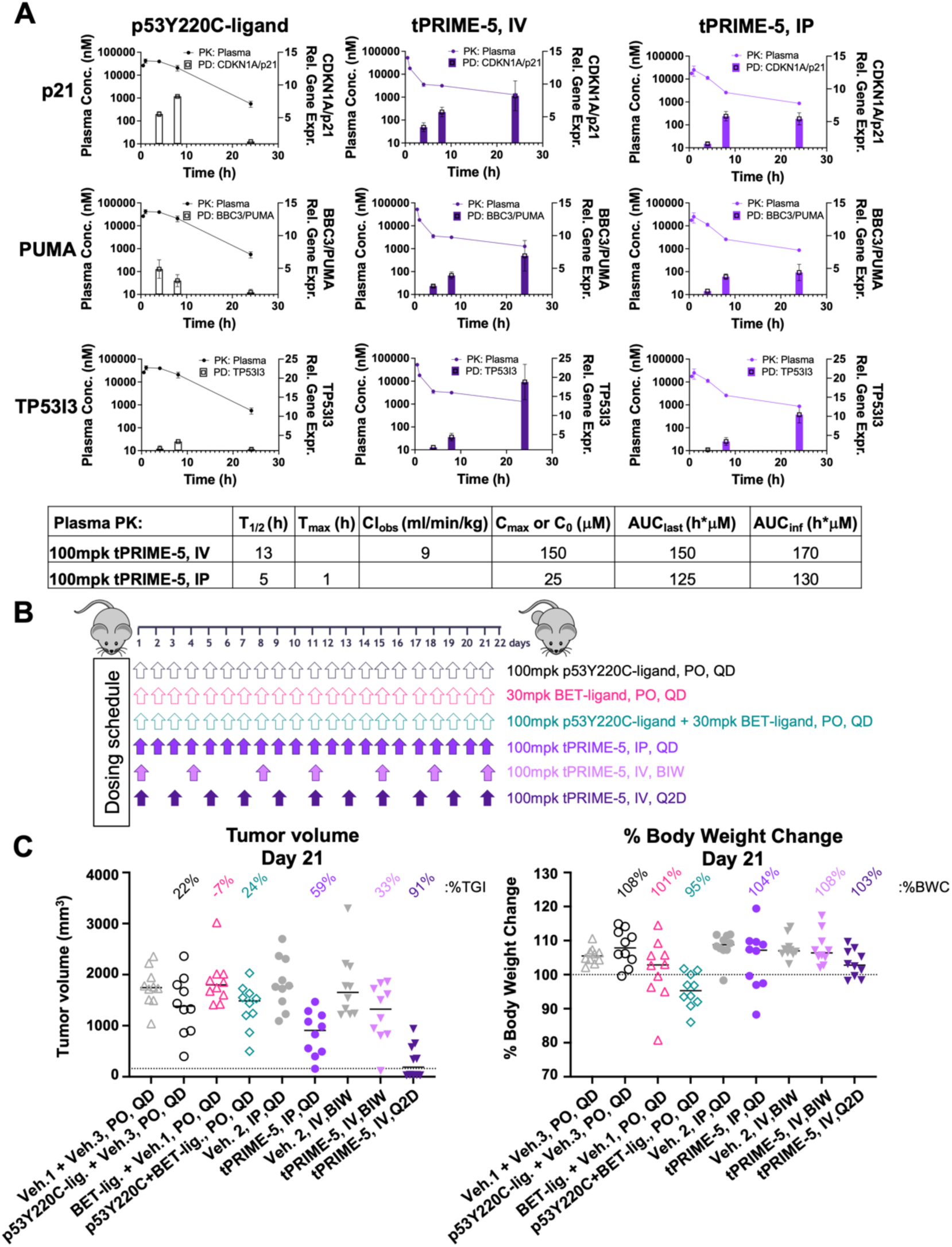
tPRIME-5 induces p53 target genes in vivo and achieves tumor growth inhibition and regression in NUGC3 xenografts. **A.** Plasma pharmacokinetics and NUGC3 tumor pharmacodynamics measured post a single dose. Left y-axis shows plasma concentration of tPRIME-5 or p53Y220C-ligand measured 0.5h, 1h, 4h, 8h or 24h post dosing mice IV, IP or PO. Right y-axis shows gene expression changes of CDKN1A, BBC3 and TP53I3 in NUGC3 xenografts measured at 4h, 8h and 24h post a single dose. **B.** Schematic of 21-day efficacy study in NUGC3 xenograft model. **C.** NUGC3 tumor growth inhibition and body weight changes of individual animals across the ten treatment groups at the end of study.

To address if tPRIME-5 can be effective in vivo, modulation of p53-target genes identified through RNA-sequencing and validated through qPCR and proteomics as PD biomarkers was assessed in NUGC3 xenografts collected 4h, 8h, and 24h post a single dose of 100mpk tPRIME-5 dosed IP or IV. tPRIME-5 achieved superior induction of p53-target genes associated with apoptosis, such as BBC3/PUMA, RRAD and TP53I3, compared to p53Y220C-ligand or p53Y220C-plus BET-ligand combined (Fig. 5A, Supplementary Fig. 9A). IV dosing resulted in more pronounced target gene induction than IP dosing, in agreement with the achieved exposures (Fig. 5A, Table). In addition to achieving a greater E_max_, tPRIME-5 induced these genes with a different kinetics compared to p53Y220C-ligand alone or combined with BET-ligand: while gene expression induced by tPRIME-5 increased from 4h to 24h and achieved highest magnitude at 24h post dose, p53Y220C-ligand alone or combined with BET-ligand showed the opposite kinetics. p53 target genes p21, GDF15, and MDM2 were induced equivalently or to a lesser degree by tPRIME-5 compared to p53Y220C-ligand treatment arms, but again with unique kinetics to the p53Y220C-ligand alone (Fig. 5, Supplementary Fig. 9). These observed trends, both the superior induction of apoptotic p53-target genes as well as the inverse kinetics, were in agreement with cellular in vitro data described above.

To test if tPRIME-5 can inhibit tumor growth in vivo, NUGC3 cells were implanted subcutaneously into the flanks of Balb/c nude mice (Fig. 5B). IV dosing of 100mpk tPRIME-5 every two days resulted in 91% tumor growth inhibition, with complete tumor regression being observed in five of the ten animals by end of study. Treatment of animals with 100mpk tPRIME-5 IV twice a week (BIW) resulted in 33% tumor growth inhibition, and daily dosing of 100mpk tPRIME-5 IP resulted in 59% tumor growth inhibition. All tPRIME-5-treatment arms were well tolerated with no significant weight loss observed. In addition, all tPRIME-5-treatment arms were more efficacious than treatment with 100mpk p53Y220C-ligand, which achieved 22% tumor growth inhibition (TGI), as well as the combination of p53Y220C-ligand + BET-ligand with 24% TGI. The combination arm was least tolerated and resulted in body weight loss over the time course of 21 days of treatment (Fig. 5C, Supplementary Fig. 10).

Pharmacodynamic biomarker changes were compared between tumor samples collected from the different treatment groups at the end of the 21-day efficacy study, with exception of the 100mpk tPRIME-5 IV Q2D treatment group, for which tumor size was too small.

Superior PD biomarker modulation was detected in tumors treated with 100mpk tPRIME-5 IP QD compared to 100mpk tPRIME-5 IV BIW, correlating with the tumor growth inhibition achieved in these groups (Supplementary Fig. 11). Both tPRIME-5 treatment groups achieved superior induction of pro-apoptotic p53 target genes ATF3, BBC3, RRAD, TP53I3 and ALDH1A3 compared to p53Y220C-ligand alone or combined with BET-ligand.

Together, these data demonstrate that tPRIME-5, even at lower dosing frequency and less favorable PK properties, achieves superior tumor growth inhibition and regression in vivo compared to p53Y220C-stabilizers due to greater induction of pro-apoptotic p53 target genes.

## Discussion

p53Y220C-stabilizers are the first small molecules that can specifically reactivate the tumor suppressor p53 by reconstituting it in its wildtype conformation. In p53Y220C-mutant cells, this mode of action results in cell proliferation inhibition, and in a Phase 1 study (NCT04585750), Rezatapopt, the first p53Y220C-stabilizer in the clinic, achieved 38% ORR at 2,000 mg QD across a range of cancers. Here, we developed heterobifunctional molecules linking p53Y220C-ligands with BET-ligands called tPRIME, which induce marked induction of apoptotic p53 target genes, potent inhibition of cell and tumor growth and apoptosis of p53Y220C mutant cancer cells in a manner uniquely superior to p53 stabilization alone.

tPRIME-5 achieved superior tumor growth inhibition compared to the p53Y220C-ligand, at lower overall exposures and less frequent dosing.

One noticeable difference between tPRIMEs and p53Y220C-ligand was the superior activation of p53-target genes associated with apoptosis rather than cell cycle arrest by tPRIMEs. It has been demonstrated that p53 response elements at cell cycle arrest genes (e.g. CDKN1A) bind p53 with higher affinity than those at apoptosis targets (e.g. BBC3)^39–41^. With this mechanism in place, lower p53 levels induced by low grade DNA damage would induce cell cycle arrest, whereas high p53 levels induced by high grade DNA damage would also induce apoptosis genes, allowing appropriate cell-fate control based on the environmental stress^41^. Interestingly, based on our DSG crosslinking and tetramer quantification data, tPRIME molecules do not induce significantly more p53 tetramer formation than p53Y220C-stabilizers, suggesting that the absolute levels of functional p53 are not solely responsible for the differential activation of apoptotic genes. It has further been proposed that changes in the DNA minor/major groove due to differences in the p53 consensus sequence can lead to distinct p53 DNA binding modes by inducing different Arg248 and Lys120 conformations and interactions^41–47^. It is possible that p53 bound by tPRIMEs in a ternary complex adopts a different conformation than p53 bound by a p53 stabilizer, enabling preferential binding to p53 consensus motifs on apoptotic p53 target genes, and may not require a tetrameric form to the same degree.

Another noticeable difference between tPRIMEs and p53Y220C-ligand was the gene activation kinetics: p53Y220C-ligands induced p53 target genes as fast as 1h post treatment followed by downregulation post 8h, correlating with the induction of MDM2, a negative regulator of p53. In contrast, tPRIMEs led to a monotonic induction of p53 target genes with gene expression increasing over the time course of 24h, while sparing, for the most part, the induction of MDM2. These differences in kinetics and MDM2-mediated feedback activation may de-regulate the transcriptional networks underlying the regulation of different p53 target genes, thereby altering the balance from cell cycle arrest to apoptosis^48^.

Another possibility for the preferential activation of apoptotic p53 target genes may be related to the BET-ligand side of the tPRIME, which may take a role in redirecting p53 to genes proximal to chromatin regions associated with BET-proteins or other chromatin interacting p-TEFb proteins. ChiP Sequencing experiments may allow elucidating this on a molecular level in the future.

Many BET-protein inhibitors have been facing challenges in the clinic due to dose-limiting toxicities and narrow therapeutic windows^13–15^, precluding their approval. One relevant question will be whether tPRIMEs, utilizing BET-ligands on one side of the molecule, will face these challenges as well. Firstly, target engagement evaluation via NanoBRET demonstrated lower binding affinity of tPRIMEs to BRD4 compared to BET-ligands in cells (BET-ligand: 0.09μM, tPRIME-5: 0.23μM; data not shown). Secondly, knockdown of all three BET-proteins BRD2, BRD3, and BRD4 was required to attenuate tPRIME-mediated apoptosis, indirectly suggesting that only a fraction of BET-proteins may be required to mediate tPRIMEs’ cellular effect, similarly to what was suggested for BCL6-BRD4-TCIPs^29^. Lastly, animals treated with tPRIME-5 did not show signs of body weight loss, whereas the combination treatment of p53Y220C-ligand and BET-ligand did. All these observations suggest that tPRIMEs may be less problematic in causing dose-limiting toxicities.

A challenge across p53Y220C-stabilizers seems to be the high drug load required to achieve a preclinical or clinical response. For example, 2,000mg of Rezatapopt was identified as the RP2D, resulting in 38% ORR in Phase 1^8^. The reason for this high drug load may lie in the nature of p53 functioning as a tetramer: in the case of the destabilizing p53Y220C-mutation, four p53 monomers need to be stabilized simultaneously to form one functional p53 transcription factor that can activate p53 target genes. It is also known that large pools of mutant p53 protein accumulate in cells carrying p53 mutations compared to p53 wild type, necessitating higher doses for full target engagement. Based on our data, tPRIMEs can achieve more pronounced p53 target gene activation and cellular phenotypes more potently and at equivalent levels of formed tetramer, suggesting that potentially lower drug load is achievable for the clinic. Another challenge for stabilizers operating primarily under cell cycle arrest / cytostatic response may be an insufficient stimulus required to overcome adaptive resistance in solid tumors. Indeed, it is notable that in the Ph1/2 trial of Rezatapopt, an exclusion criterion is the co-occurrence of KRAS mutations. Notably p53Y220C-BET-tPRIMEs demonstrate potent activity in cell lines independent of KRAS status (e.g., T3M4, KRAS Q61H, Supplementary Fig. 1A).

Lastly, given that p53 mutations lead to protein inactivation, thereby alleviating the negative feedback loop, p53 mutant cells, including p53Y220C cell lines express high levels of p53. p53-PLK1-heterobifunctionals take advantage of this phenomenon by p53-mediated accumulation of compound in the cells and selectively inhibiting p53Y220C-mutant cell growth through inhibiting the essential protein PLK1^28^. This RIPTAC concept has also been put forward by Halda Therapeutics, demonstrating killing of androgen receptor expressing prostate cells though inhibition of BRD4 as essential protein^49^. For p53Y220C-BET-tPRIMEs, we could not see evidence of a RIPTAC-like mechanism but instead demonstrated strong activation of p53Y220C and its target genes. It will be interesting to better understand and potentially predict in the future the rules determining effector or recruiter based on the cellular target or context. The p53Y220C-BET-tPRIMEs described herein have since been further optimized to achieve enhanced potency and improved oral exposure. The results of these ongoing efforts will be reported in due course.

In summary, our data on p53Y220C-tPRIMES suggest that these molecules provide a compelling alternative as a therapeutic modality compared to p53Y220C-stabilizers for p53Y220C mutant cancers.

---

**Supplementary Figure 1:**
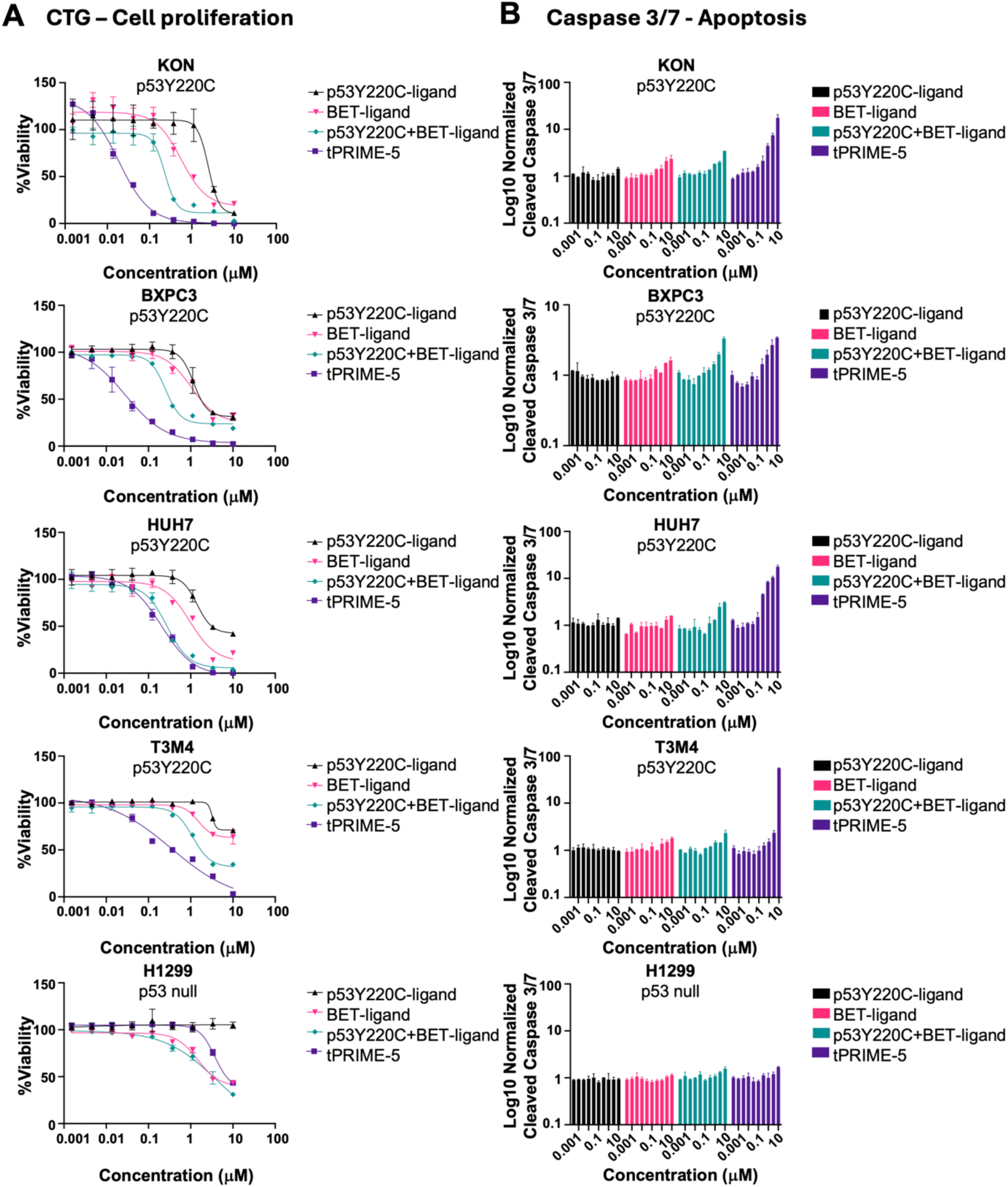
tPRIME-5 inhibits cell proliferation and induces apoptosis across a panel of p53Y220C-mutant cell lines. **A.** Cell viability assessment via CTG after 96h treatment of p53Y220C-mutant cell lines KON, BXPC3, HUH7 and T3M4 or p53 null cell line H1299 with tPRIME-5, p53Y220C-ligand, BET-ligand or combination of p53Y220C+BET-ligand. **B.** Apoptosis assessment via cleaved caspase 3/7 after 24h treatment of p53Y220C-mutant cell lines KON, BXPC3, HUH7 and T3M4 or p53 null cell line H1299 with tPRIME-5, p53Y220C-ligand, BET-ligand or combination of p53Y220C+BET-ligand.

**Supplementary Figure 2:**
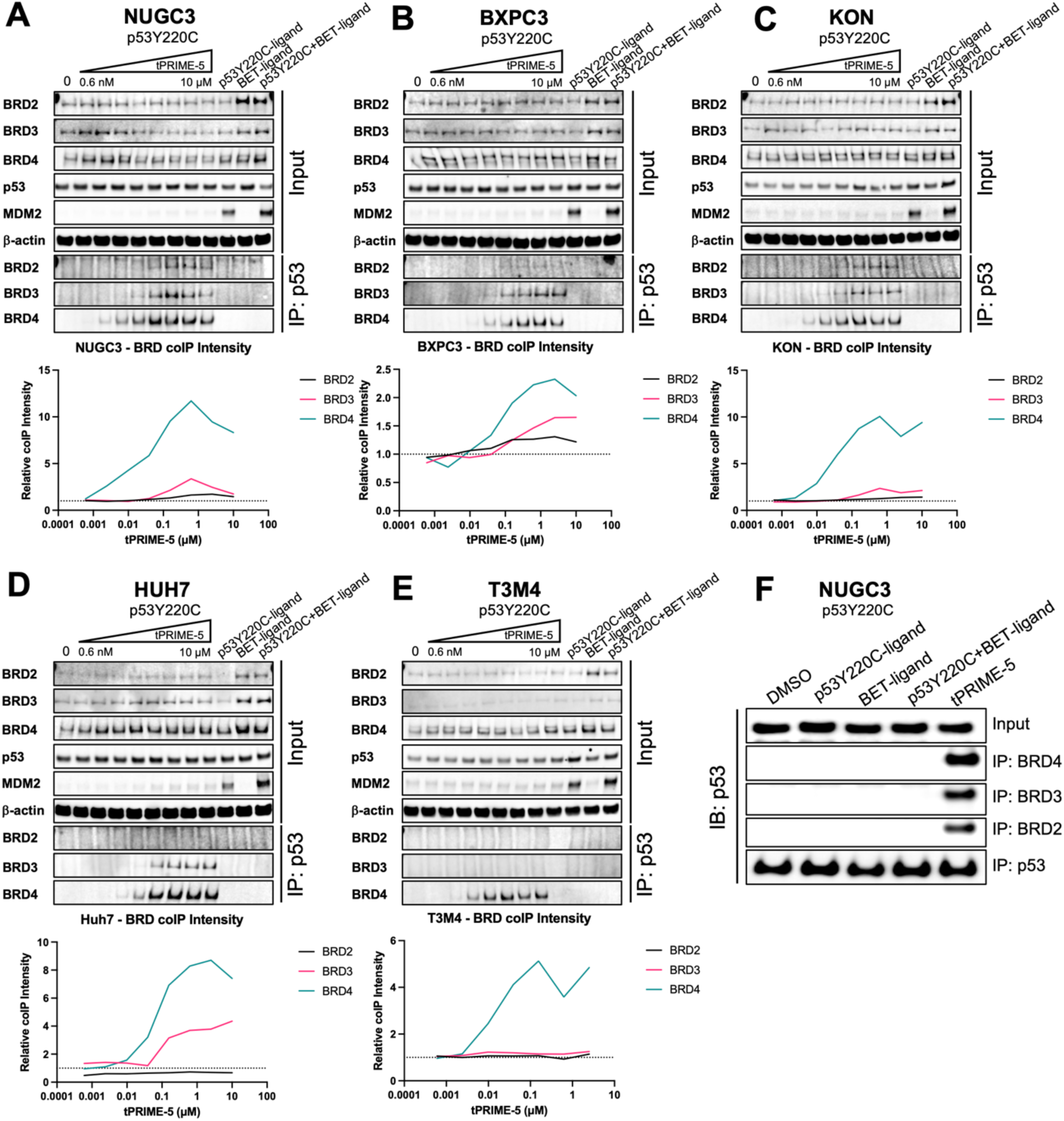
tPRIME-5 induces ternary complex formation between p53Y220C and BET-proteins across a panel of p53Y220C-mutant cell lines. **A-E.** Western blot analysis demonstrating co-immunoprecipitation of BRD2, BRD3 and BRD4 after pulldown of p53 across multiple p53Y220C-mutant cell lines treated with serially diluted tPRIME-5 for 4h. **F.** Western Blot analysis demonstrating co-immunoprecipitation of p53 after pulldown of BRD2, BRD3, or BRD4 in NUGC3 cells treated with 1μM tPRIME-5 or control ligands for 4h.

**Supplementary Figure 3:**
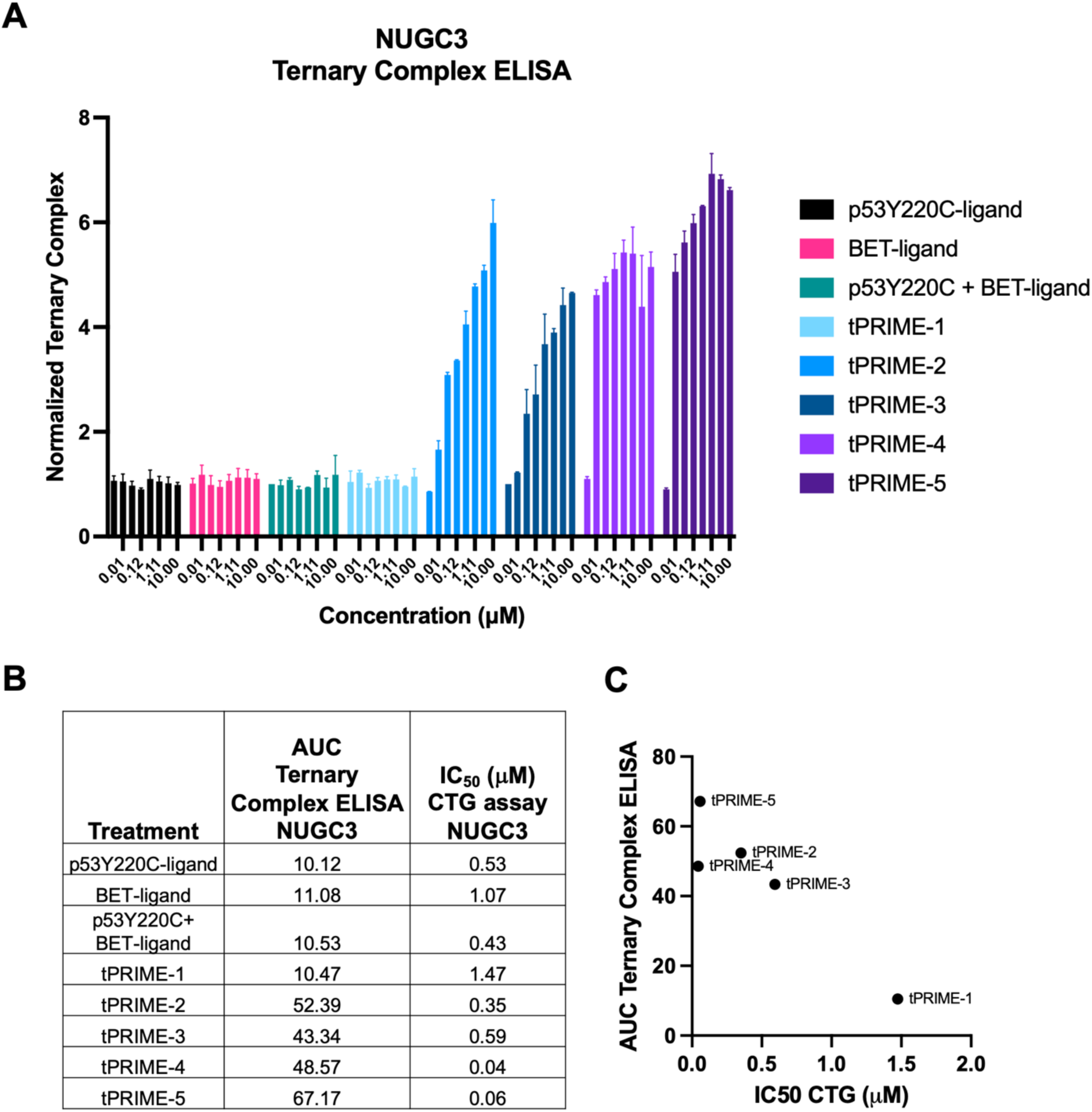
Ternary complex formation correlates with efficacy in CTG proliferation assay. **A.** Assessment of ternary complex formation via ELISA in NUGC3 cells treated with serially diluted tPRIME-1-5 and control ligands for 4h. **B.** AUC values from ternary complex ELISA assay in NUGC3 cells post 4h treatment and IC_50_ (μM) values from 96h CTG cell proliferation assay in NUGC3. **C.** Scatter plot of AUC values from NUGC3 ternary complex ELISA versus IC50 values from NUGC3 CTG cell proliferation assay.

**Supplementary Figure 4:**
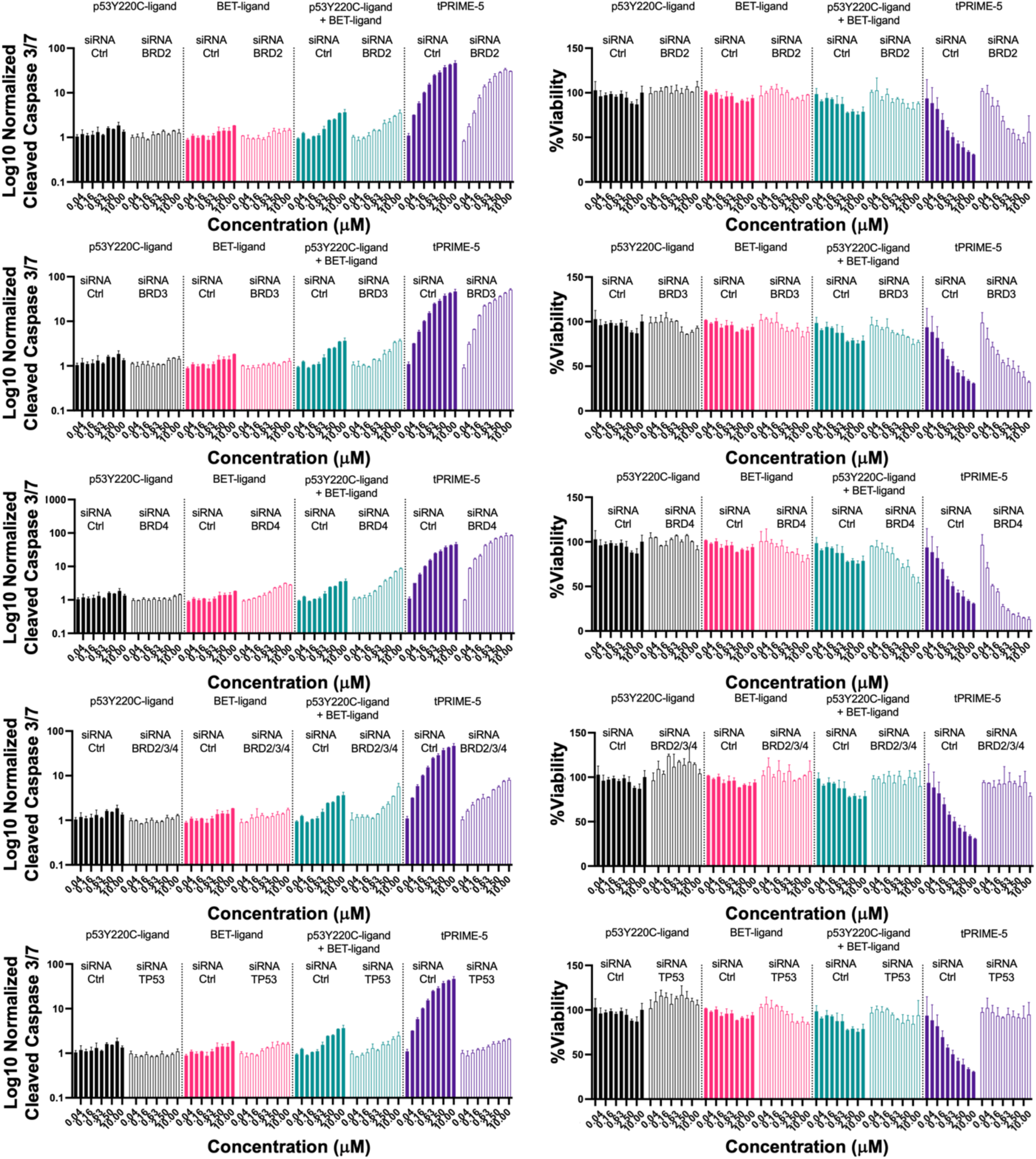
Knockdown of TP53 or combined knockdown of BRD2/3/4, but not individual knockdown of BRD2, BRD3, or BRD4, attenuates tPRIME-5-induced apoptosis and cell viability reduction. Caspase 3/7 and CTG cell viability were assessed 24h post treatment of NUGC3 cells with serially diluted tPRIME-5 or control ligands in the presence of TP53 siRNA knockdown, combined knockdown of BRD2, BRD3 and BRD4 or individual knockdown of BRD2, BRD3 or BRD4.

**Supplementary Figure 5:**
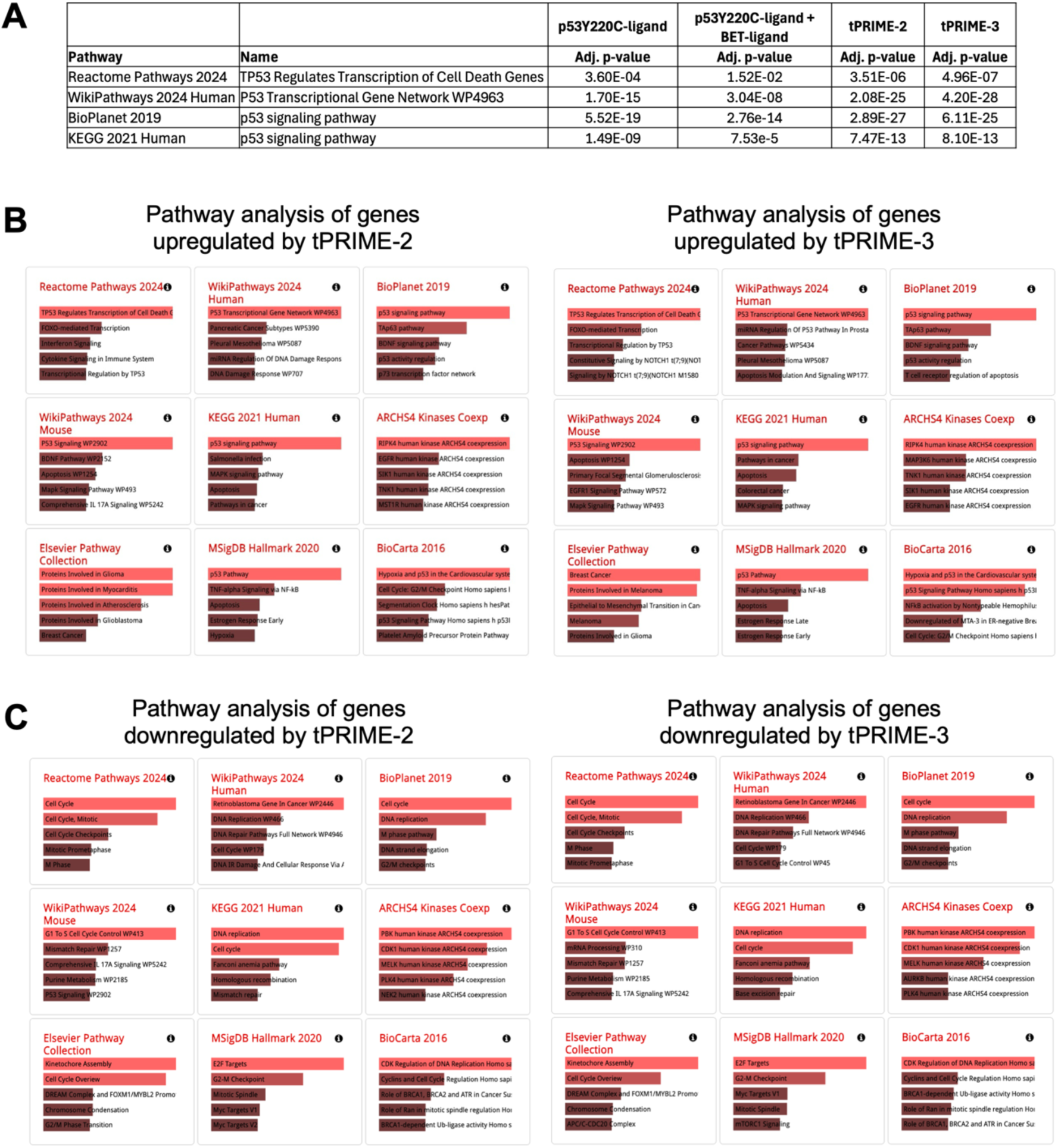
Pathway analysis of genes significantly up- or downregulated in RNA-sequencing experiment in NUGC3 cells treated with tPRIMEs or control ligands compared to DMSO. **A.** ENRICHR pathway analysis of genes upregulated by different treatments (1μM p53Y220C-ligand, 1μM BET-ligand, 1μM p53Y220C-ligand + 1μM BET-ligand, 1μM tPRIME-2, 1μM tPRIME-3) compared to DMSO: p-value ≤ 0.05; Log2 mean expression treatment ≥ 2; Log2 Fold expression compared to DMSO ≥ 1. **B.** Images of results from ENRICHR pathway analysis of genes upregulated by tPRIME-2/3 compared to DMSO (p-value ≤ 0.05; Log2 mean expression treatment ≥ 2; Log2 Fold expression compared to DMSO ≥ 1). **C.** Images of results from ENRICHR pathway analysis of genes downregulated by tPRIME-2/3 compared to DMSO (p-value ≤ 0.05; Log2 mean expression DMSO ≥ 2; Log2 Fold expression compared to DMSO ≤ 1).

**Supplementary Figure 6:**
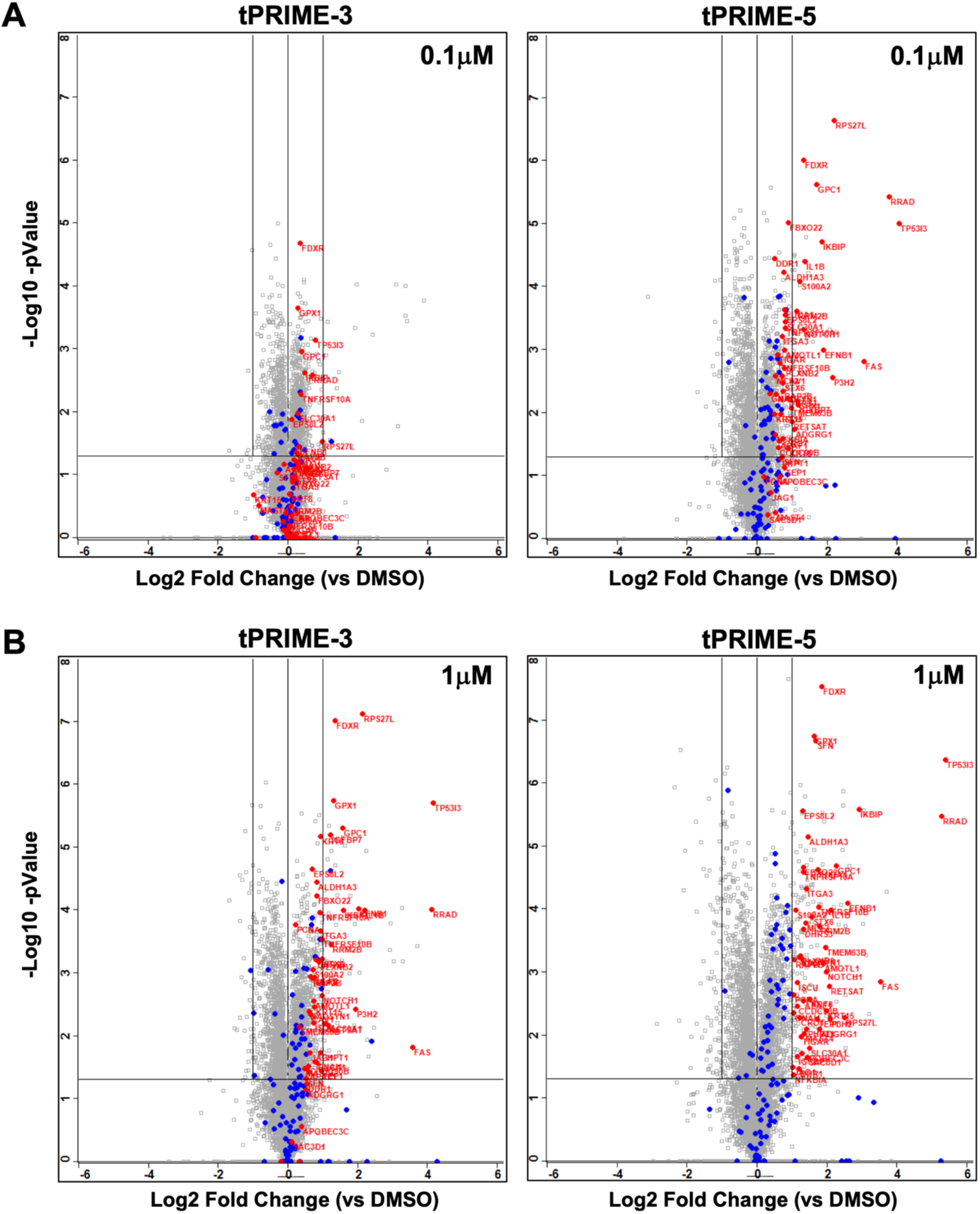
tPRIME-3 and tPRIME-5 significantly induce p53-dependent target proteins. Volcano plot of proteins identified through LC-MS/MS in NUGC3 cells treated for 24h with 0.1μM (**A**) or 1μM (**B**) of tPRIME-3 or tPRIME-5 relative to DMSO.

**Supplementary Figure 7:**
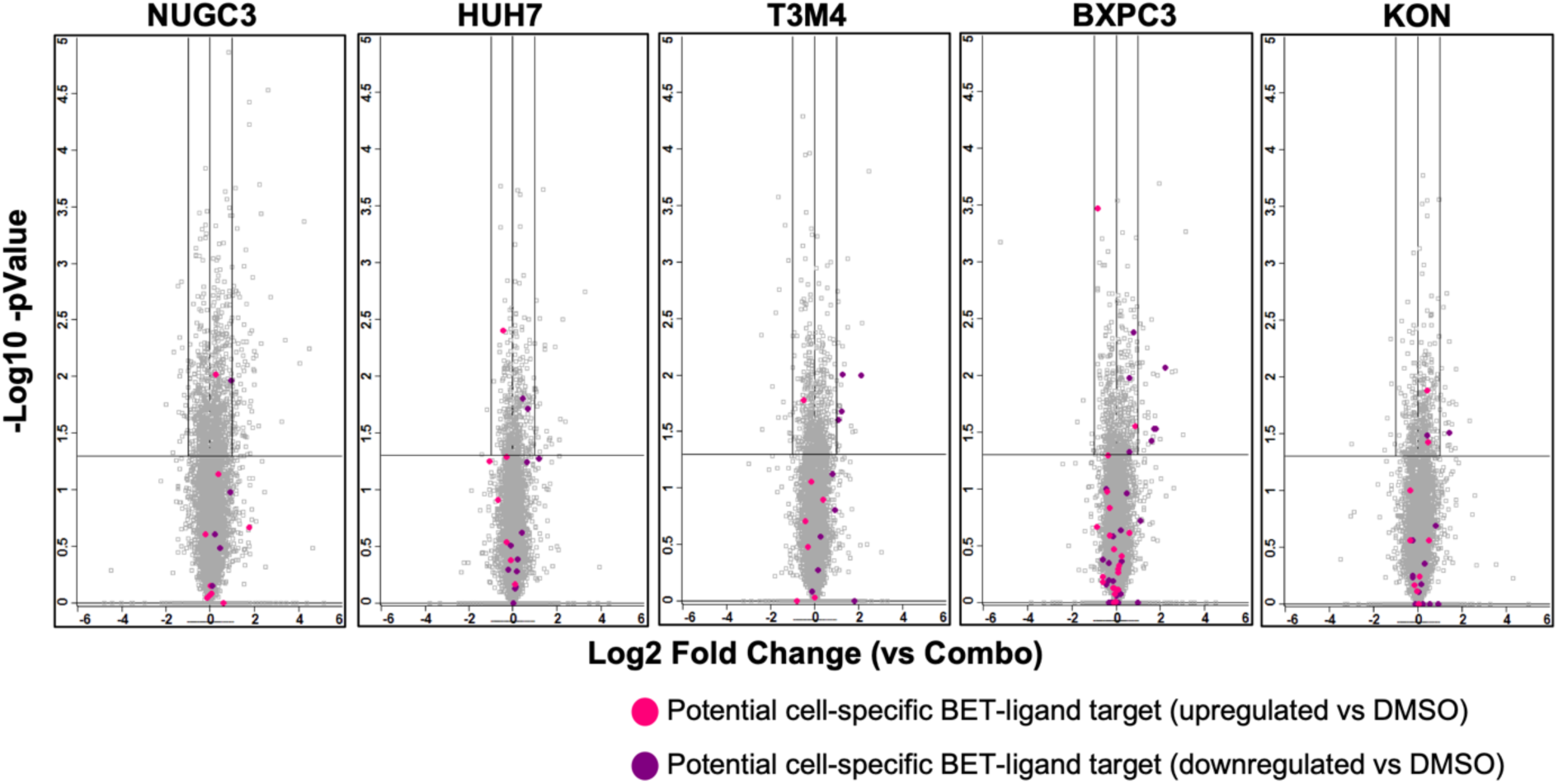
tPRIME-5 does not meaningfully impact cell-specific BET target genes. Volcano plot of proteins identified through LC-MS/MS in several p53Y220C-mutant cells treated for 24h with 1μM of tPRIME-5 compared to combination of p53Y220C+BET-ligand combination.

**Supplementary Figure 8:**
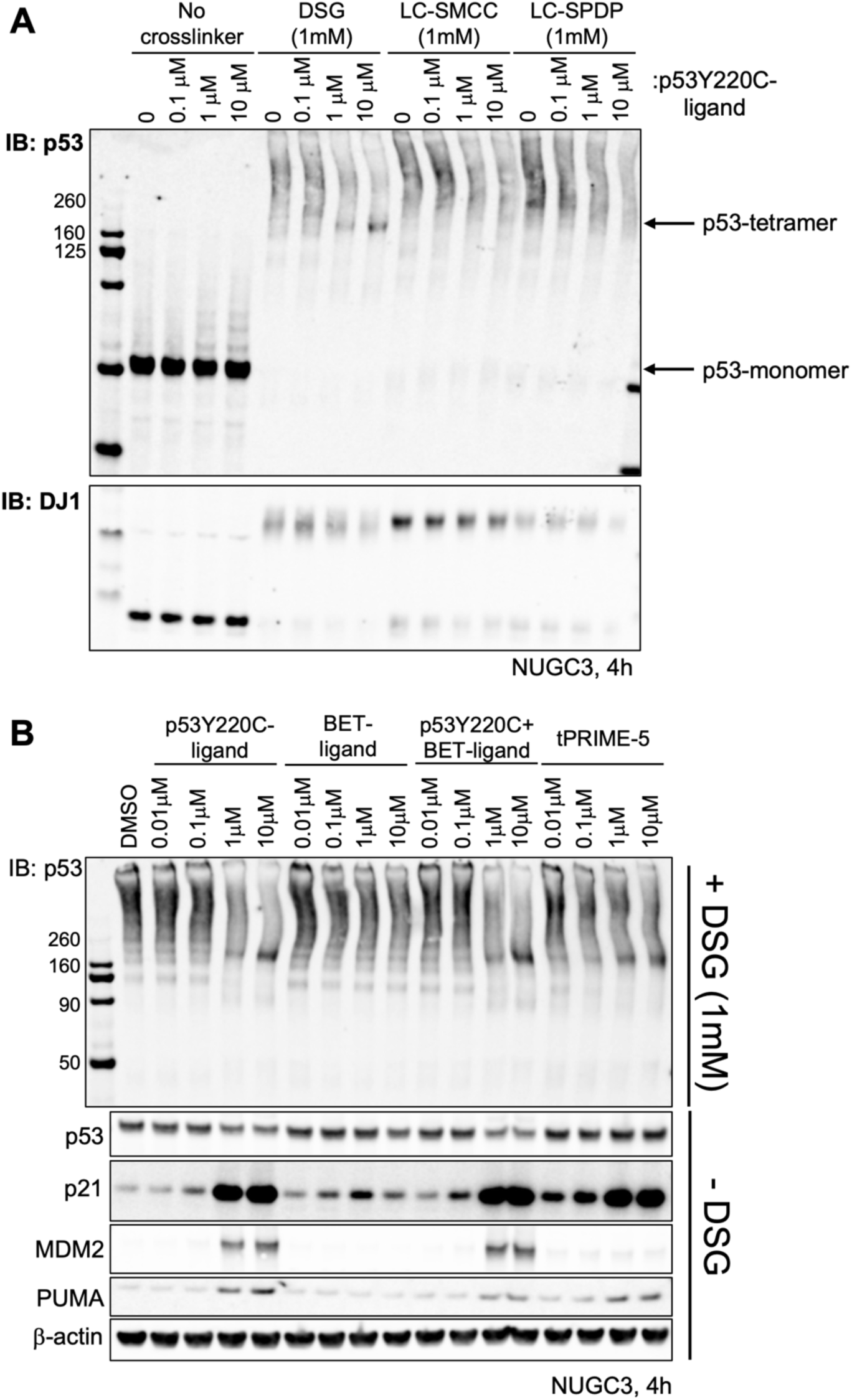
tPRIME-5 induces p53 tetramer formation at a lower concentration than p53Y220C-ligand. **A.** DSG crosslinker, but not LC-SMCC or LC-SPDP crosslinker, allows the detection of p53 tetramer in NUGC3 cells post 4h treatment with 1μM or 10μM p53Y220C-ligand. **B.** p53Y220C-ligand and tPRIME-5 induce p53-tetramer formation in NUGC3 cells dose-dependently post 4h treatment.

**Supplementary Figure 9:**
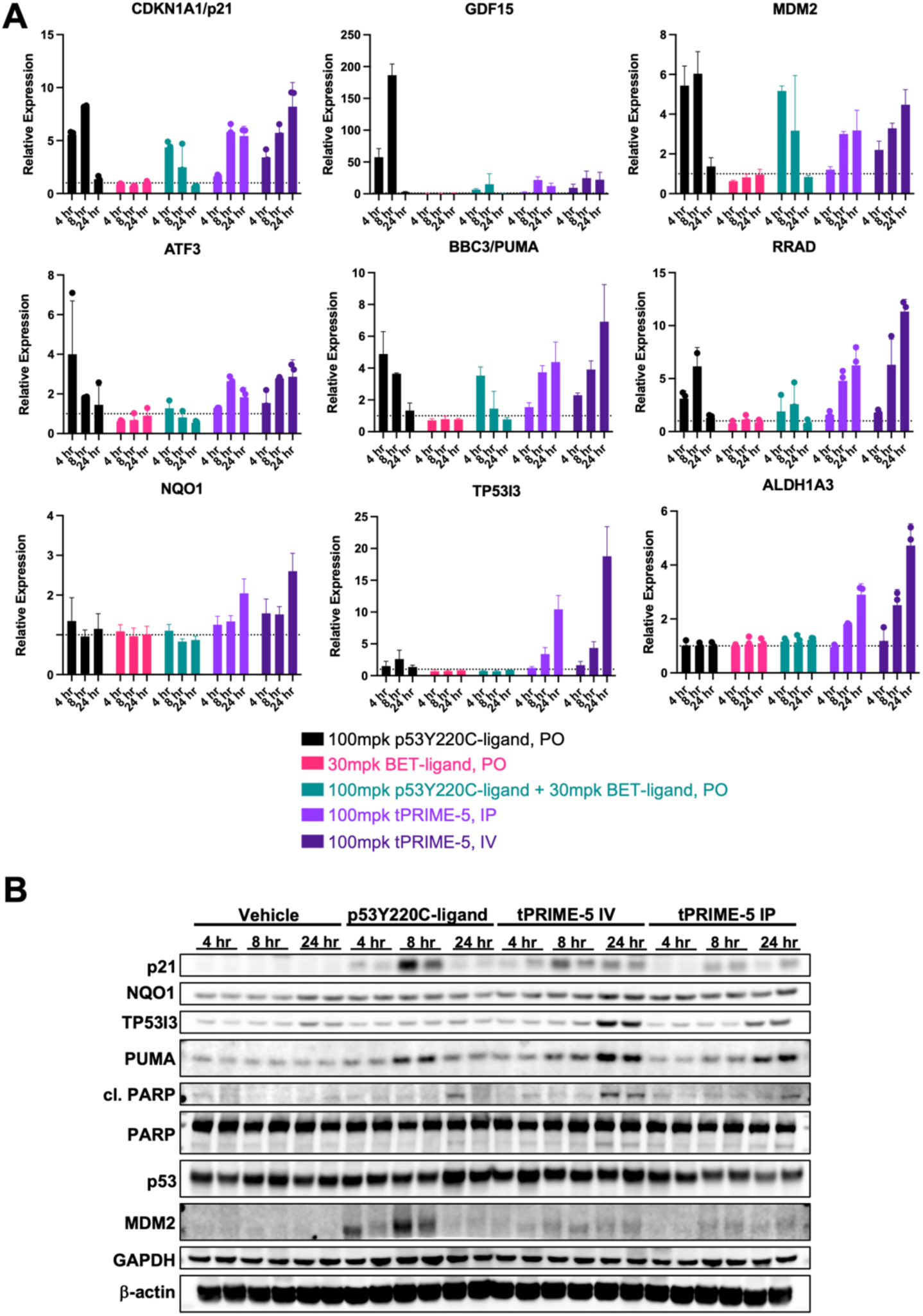
tPRIME-5 induces marked expression of multiple p53 target genes in NUGC3 xenografts post a single dose. **A.** qPCR analysis of the p53 target genes CDKN1A, GDF15, MDM2, ATF3, BBC3, RRAD, NQO1, TP53I3 and ALDH1A3 in NUGC3 xenografts treated with a single dose of vehicle, 100mpk p53Y220C-ligand (PO), 30mpk BET-ligand (PO), 100mpk p53Y220C-ligand + 30mpk BET-ligand (PO), 100mpk tPRIME-5 (IV) or 100mpk tPRIME-5 (IP) and harvested 4h, 8h and 24h post dosing. **B.** Analysis of p53 dependent target proteins via Western Blot in NUGC3 xenograft samples harvested 4h, 8h and 24h post a single dose of treatment with vehicle, 100mpk p53Y220C-ligand (PO) or 100mpk tPRIME-5 (IV or IP).

**Supplementary Figure 10:**
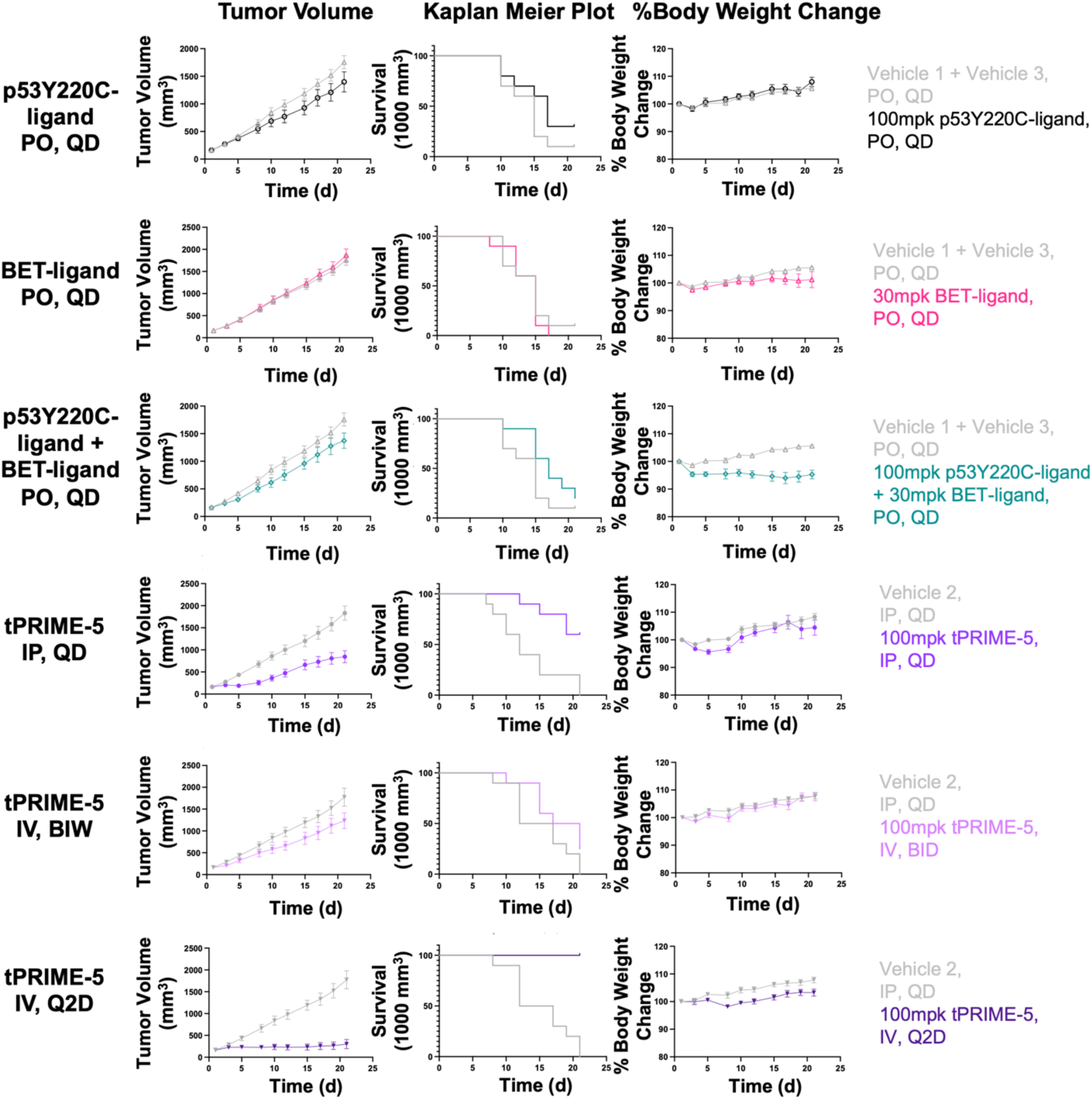
tPRIME-5 achieves greater tumor growth inhibition and survival benefit without significant body weight loss in NUGC3 xenografts compared to p53Y220C-ligand alone or combined with BET-ligand. Tumor volume (mm^3^), Kaplan Meier Plot and %Body Weight change over time shown for the different treatment arms. For Kaplan Meier Plot, tumor volume exceeding 1,000mm^3^ is categorized as survival loss.

**Supplementary Figure 11:**
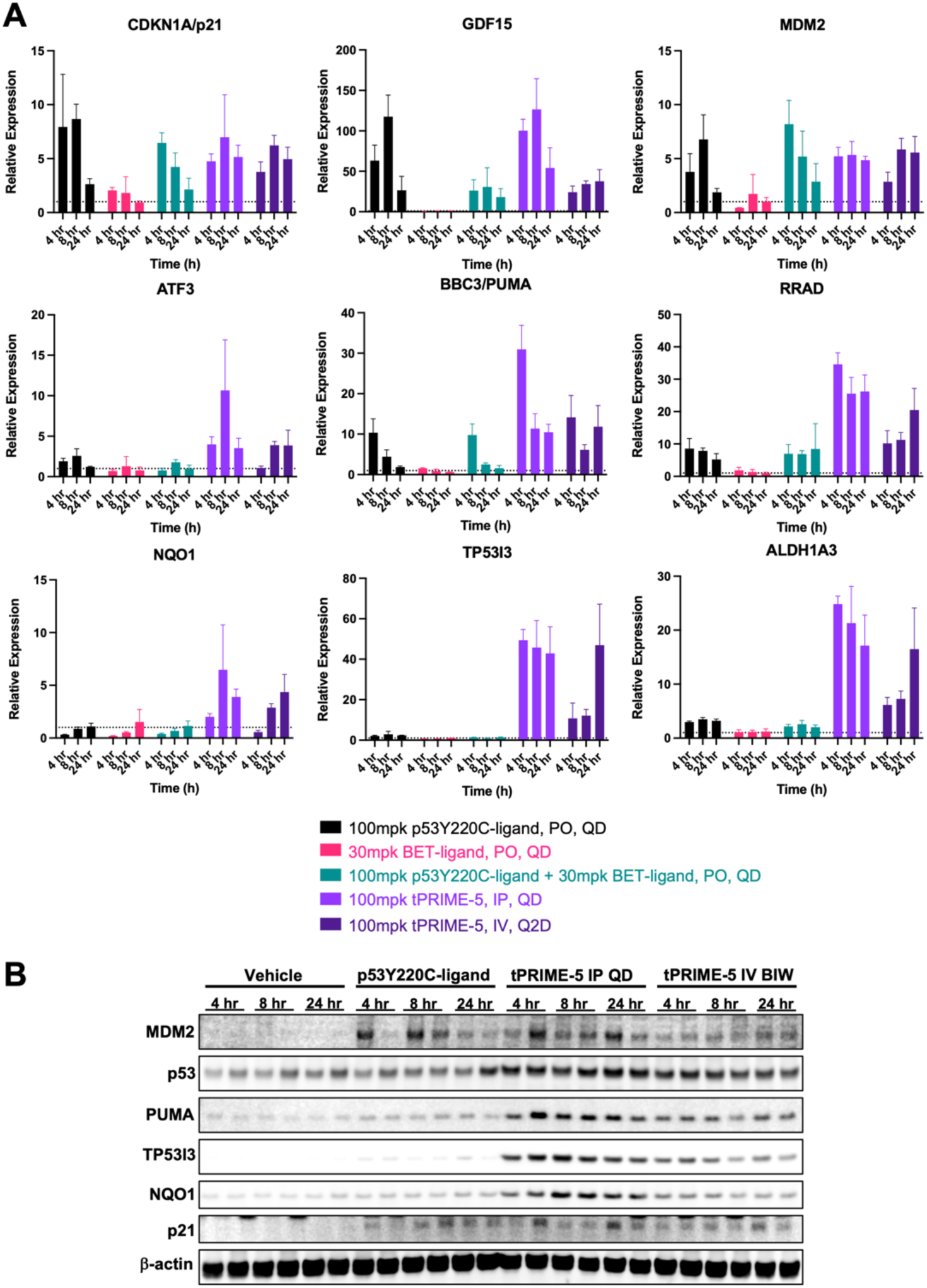
tPRIME-5 induces marked expression of multiple p53 target genes in NUGC3 xenografts harvested at the end of the efficacy study. **A.** qPCR analysis of the p53 target genes CDKN1A, GDF15, MDM2, ATF3, BBC3, RRAD, NQO1, TP53I3 and ALDH1A3 in NUGC3 xenografts treated for 21 days with vehicle, 100mpk p53Y220C-ligand (PO QD), 30mpk BET-ligand (PO QD), 100mpk p53Y220C-ligand + 30mpk BET-ligand (PO QD), 100mpk tPRIME-5 (IV BIW) or 100mpk tPRIME-5 (IP QD) and harvested 4h, 8h and 24h post last dose. **B.** Analysis of p53 dependent target proteins via Western Blot in NUGC3 xenograft samples harvested 4h, 8h and 24h post last treatment with vehicle, 100mpk p53Y220C-ligand (PO QD) or 100mpk tPRIME-5 (IV BIW or IP QD).

**Supplementary Table 1:** IC_50_ values (μM) from 96h CTG cell proliferation assay across different cell lines.

|  | **GeoMean IC<sub>50</sub> (μM)** |  |  |  |  |  |  |
| --- | --- | --- | --- | --- | --- | --- | --- |
|  | **NUGC3**<br>**p53Y220C** | **BXPC3**<br>**p53Y220C** | **HUH7**<br>**p53Y220C** | **KON**<br>**p53Y220C** | **T3M4**<br>**p53Y220C** | **HCT116**<br>**p53 wt** | **H1299**<br>**p53 null** |
| **p53Y220C-ligand** | 0.787 | 0.559 | 0.450 | 1.290 | > 4.69 | > 9.30 | > 9.99 |
| **BET-ligand** | 1.070 | 1.020 | 0.958 | 0.557 | 1.510 | > 10.0 | 1.460 |
| **p53Y220C+BET-ligand** | 0.413 | 0.253 | 0.274 | 0.230 | 1.120 | > 10.0 | 4.010 |
| **tPRIME-1** | 1.730 | 0.438 | 3.720 | 0.411 | 2.610 | 3.320 | 6.830 |
| **tPRIME-2** | 0.333 | 0.269 | 0.969 | 0.204 | > 1.04 | > 5.62 | > 4.84 |
| **tPRIME-3** | 0.601 | 0.138 | 0.472 | 0.174 | 0.891 | > 8.26 | > 7.45 |
| **tPRIME-4** | 0.044 | 0.015 | 0.047 | 0.013 | 0.2418 | 4.480 | 1.240 |
| **tPRIME-5** | 0.046 | 0.018 | 0.099 | 0.034 | 0.165 | > 5.49 | 4.140 |

## References

1. Vousden, K. H. & Lane, D. P. p53 in health and disease. Nat. Rev. Mol. Cell Biol. 8, 275– 283 (2007).

2. Boeckler, F. M. et al. Targeted rescue of a destabilized mutant of p53 by an in silico screened drug. Proc. Natl. Acad. Sci. U. S. A. 105, 10360–10365 (2008).

3. Joerger, A. C., Ang, H. C. & Fersht, A. R. Structural basis for understanding oncogenic p53 mutations and designing rescue drugs. Proc. Natl. Acad. Sci. U. S. A. 103, 15056– 15061 (2006).

4. Fallatah, M. M. J., Law, F. V., Chow, W. A. & Kaiser, P. Small-molecule correctors and stabilizers to target p53. Trends Pharmacol. Sci. 44, 274–289 (2023).

5. Guiley, K. Z. & Shokat, K. M. A Small Molecule Reacts with the p53 Somatic Mutant Y220C to Rescue Wild-type Thermal Stability. Cancer Discov. 13, 56–69 (2023).

6. Puzio-Kuter, A. M. et al. Restoration of the Tumor Suppressor Function of Y220C-Mutant p53 by Rezatapopt, a Small-Molecule Reactivator. Cancer Discov. 15, 1159–1179 (2025).

7. Vu, B. T. et al. Discovery of Rezatapopt (PC14586), a First-in-Class, Small-Molecule Reactivator of p53 Y220C Mutant in Development. ACS Med. Chem. Lett. 16, 34–39 (2025).

8. Schram, A. M., et al. Abstract LB_A25: Updated Phase 1 results from the PYNNACLE Phase 1/2 study of PC14586, a selective p53 reactivator, in patients with advanced solid tumors harboring a TP53 Y220C mutation. Mol. Cancer Ther. 22, LB_A25 (2023).

9. Belkina, A. C. & Denis, G. V. BET domain co-regulators in obesity, inflammation and cancer. Nat. Rev. Cancer 12, 465–477 (2012).

10. Nicodeme, E. et al. Suppression of inflammation by a synthetic histone mimic. Nature 468, 1119–1123 (2010).

11. Jang, M. K. et al. The bromodomain protein Brd4 is a positive regulatory component of P-TEFb and stimulates RNA polymerase II-dependent transcription. Mol. Cell 19, 523– 534 (2005).

12. Pérez-Salvia, M. & Esteller, M. Bromodomain inhibitors and cancer therapy: From structures to applications. Epigenetics 12, 323–339 (2016).

13. Trojer, P. Targeting BET Bromodomains in Cancer. Annu. Rev. Cancer Biol. 6, 313– 336 (2022).

14. Shorstova, T., Foulkes, W. D. & Witcher, M. Achieving clinical success with BET inhibitors as anti-cancer agents. Br. J. Cancer 124, 1478–1490 (2021).

15. Sun, Y. et al. Safety and Efficacy of Bromodomain and Extra-Terminal Inhibitors for the Treatment of Hematological Malignancies and Solid Tumors: A Systematic Study of Clinical Trials. Front. Pharmacol. 11, 621093 (2020).

16. Singh, S. et al. Proximity-inducing modalities: the past, present, and future. Chem. Soc. Rev. 52, 5485–5515 (2023).

17. Nalawansha, D. A., Mangano, K., den Besten, W. & Potts, P. R. TAC-tics for Leveraging Proximity Biology in Drug Discovery. *Chembiochem Eur*. J. Chem. Biol. 25, e202300712 (2024).

18. Sakamoto, K. M. et al. Protacs: Chimeric molecules that target proteins to the Skp1–Cullin–F box complex for ubiquitination and degradation. Proc. Natl. Acad. Sci. 98, 8554–8559 (2001).

19. Rutherford, K. A. & McManus, K. J. PROTACs: Current and Future Potential as a Precision Medicine Strategy to Combat Cancer. Mol. Cancer Ther. 23, 454–463 (2024).

20. Pergu, R. et al. Phosphorylation-Inducing Molecules for Regulating Dynamic Cellular Processes. J. Am. Chem. Soc. 147, 25316–25324 (2025).

21. Siriwardena, S. U. et al. Phosphorylation-Inducing Chimeric Small Molecules. J. Am. Chem. Soc. 142, 14052–14057 (2020).

22. Henning, N. J. et al. Deubiquitinase-targeting chimeras for targeted protein stabilization. Nat. Chem. Biol. 18, 412–421 (2022).

23. Hu, X. et al. Design, Synthesis, and Evaluation of p53Y220C Acetylation Targeting Chimeras (AceTACs). J. Med. Chem. 67, 14633–14648 (2024).

24. Kabir, M. et al. Harnessing the TAF1 Acetyltransferase for Targeted Acetylation of the Tumor Suppressor p53. Adv. Sci. Weinh. Baden-Wurtt. Ger. 12, e2413377 (2025).

25. Ng, C. S. C., Liu, A., Cui, B. & Banik, S. M. Targeted protein relocalization via protein transport coupling. Nature 633, 941–951 (2024).

26. Gibson, W. J. et al. Bifunctional Small Molecules That Induce Nuclear Localization and Targeted Transcriptional Regulation. J. Am. Chem. Soc. 145, 26028–26037 (2023).

27. Raina, K. et al. Regulated induced proximity targeting chimeras-RIPTACs-A heterobifunctional small molecule strategy for cancer selective therapies. Cell Chem. Biol. 31, 1490–1502.e42 (2024).

28. Sadagopan, A. et al. p53 protein abundance is a therapeutic window across TP53 mutant cancers and is targetable with proximity inducing small molecules. 2024.07.27.605429 Preprint at 10.1101/2024.07.27.605429 (2024).

29. Gourisankar, S. et al. Rewiring cancer drivers to activate apoptosis. Nature 620, 417–425 (2023).

30. Sarott, R. C. et al. Relocalizing transcriptional kinases to activate apoptosis. Science 386, eadl5361 (2024).

31. Zhu, X. et al. Activating p53Y220C with a Mutant-Specific Small Molecule. *bioRxiv* 2024.10.23.619961 (2024) doi:10.1101/2024.10.23.619961.

32. Wu, S.-Y., Lee, A.-Y., Lai, H.-T., Zhang, H. & Chiang, C.-M. Phospho switch triggers Brd4 chromatin binding and activator recruitment for gene-specific targeting. Mol. Cell 49, 843–857 (2013).

33. Stewart, H. J. S., Horne, G. A., Bastow, S. & Chevassut, T. J. T. BRD4 associates with p53 in DNMT3A-mutated leukemia cells and is implicated in apoptosis by the bromodomain inhibitor JQ1. Cancer Med. 2, 826–835 (2013).

34. Fischer, M., Quaas, M., Steiner, L. & Engeland, K. The p53-p21-DREAM-CDE/CHR pathway regulates G2/M cell cycle genes. Nucleic Acids Res. 44, 164–174 (2016).

35. Fischer, M. Census and evaluation of p53 target genes. Oncogene 36, 3943–3956 (2017).

36. Momand, J., Zambetti, G. P., Olson, D. C., George, D. & Levine, A. J. The mdm-2 oncogene product forms a complex with the p53 protein and inhibits p53-mediated transactivation. Cell 69, 1237–1245 (1992).

37. Wu, X., Bayle, J. H., Olson, D. & Levine, A. J. The p53-mdm-2 autoregulatory feedback loop. Genes Dev. 7, 1126–1132 (1993).

38. Dettmer, U., Newman, A. J., Luth, E. S., Bartels, T. & Selkoe, D. In vivo cross-linking reveals principally oligomeric forms of α-synuclein and β-synuclein in neurons and non-neural cells. J. Biol. Chem. 288, 6371–6385 (2013).

39. Inga, A., Storici, F., Darden, T. A. & Resnick, M. A. Differential Transactivation by the p53 Transcription Factor Is Highly Dependent on p53 Level and Promoter Target Sequence. Mol. Cell. Biol. 22, 8612–8625 (2002).

40. Szak, S. T., Mays, D. & Pietenpol, J. A. Kinetics of p53 binding to promoter sites in vivo. Mol. Cell. Biol. 21, 3375–3386 (2001).

41. Farkas, M. et al. Distinct mechanisms control genome recognition by p53 at its target genes linked to different cell fates. Nat. Commun. 12, 484 (2021).

42. Kitayner, M. et al. Structural basis of DNA recognition by p53 tetramers. Mol. Cell 22, 741–753 (2006).

43. Petty, T. J. et al. An induced fit mechanism regulates p53 DNA binding kinetics to confer sequence specificity. EMBO J. 30, 2167–2176 (2011).

44. Lukman, S., Lane, D. P. & Verma, C. S. Mapping the Structural and Dynamical Features of Multiple p53 DNA Binding Domains: Insights into Loop 1 Intrinsic Dynamics. PLOS ONE 8, e80221 (2013).

45. Pan, Y. & Nussinov, R. Lysine120 interactions with p53 response elements can allosterically direct p53 organization. PLoS Comput. Biol. 6, e1000878 (2010).

46. Tang, Y., Luo, J., Zhang, W. & Gu, W. Tip60-dependent acetylation of p53 modulates the decision between cell-cycle arrest and apoptosis. Mol. Cell 24, 827–839 (2006).

47. Sykes, S. M. et al. Acetylation of the p53 DNA-binding domain regulates apoptosis induction. Mol. Cell 24, 841–851 (2006).

48. Dean, J. A. et al. Functional consequences of a p53-MDM2-p21 incoherent feedforward loop. BioRxiv Prepr. Serv. Biol. 2024.06.25.600070 (2024) doi:10.1101/2024.06.25.600070.

49. Mullard, A. Induced proximity pushes beyond protein degraders, as first RIPTAC moves into the clinic. Nat. Rev. Drug Discov. 24, 235–237 (2025).

